# Pre-piRNA trimming safeguards piRNAs against erroneous targeting by RNA-dependent RNA Polymerase

**DOI:** 10.1101/2023.09.26.559619

**Authors:** Benjamin Pastore, Hannah L. Hertz, Wen Tang

## Abstract

In animal germ lines, The Piwi/piRNA pathway plays a crucial role in safeguarding genome integrity and promoting fertility. Following transcription from discrete genomic loci, piRNA precursors undergo nucleolytic processing at both 5’ and 3’ ends. The ribonuclease PARN-1 and its orthologs mediate piRNA 3’ trimming in worms, insects and mammals. Yet, the significance of this evolutionarily conserved processing step is not well understood. Employing *C. elegans* as a model organism, our recent work has demonstrated that 3’ trimming protects piRNAs against non-templated nucleotide additions and degradation. In this study, we present an unexpected finding that *C. elegans* deficient for PARN-1 accumulate a heretofore uncharacterized RNA species termed anti-piRNAs, which are antisense to piRNAs. These anti-piRNAs associate with Piwi proteins and display the propensity for a length of 17-19 nucleotides and 5’ guanine and adenine residues. We show that untrimmed pre-piRNAs in *parn-1* mutants are modified by the terminal nucleotidyltransferase RDE-3 and erroneously targeted by the RNA-dependent RNA polymerase EGO-1, thereby giving rise to anti-piRNAs. Taken together, our work identifies a previously unknown class of small RNAs upon loss of *parn-1* and provides mechanistic insight to activities of RDE-3, EGO-1 and Piwi proteins.

## INTRODUCTION

The Piwi protein, a germline-enriched Argonaute, and Piwi-interacting RNAs (piRNAs) are essential for gametogenesis and fertility in all animals studied to date (1-4). The primary role of the piRNA pathway is to safeguard the integrity of the germline genome by suppressing foreign sequences such as transposable elements and viral genomes (5-9). Piwi/piRNA complexes, via base-pairing interaction, target foreign transcripts and trigger epigenetic silencing (10,11). While piRNAs vary in genomic origins, lengths, and sequences across species, the mechanisms underlying their maturation are somewhat conserved (1-4). In organisms ranging from worms, insects, and to mammals, single-stranded precursor transcripts are produced from piRNA genes. These precursors are then transported to perinuclear nuage, germ granules, or the surface of mitochondria, where they undergo extensive nucleolytic processing to yield mature piRNAs with specific lengths (1-4). The final step in piRNA maturation is the 2’–*O*–methylation at their 3’ termini, catalyzed by a conserved methyltransferase (12-14).

With powerful genetic and molecular toolkit, *C. elegans* has emerged as one of the crucial organisms for studying the piRNA pathway. *C. elegans* express a single functional Piwi protein known as PRG-1. Mature piRNAs are commonly referred to as 21U-RNAs due to their strong propensity for a 5’ uridine (U) and a primary length of 21-nucleotides (nts) (15-19). Type I and II piRNAs are found in *C. elegans* and its sibling species *C. briggsae*, with Type I piRNAs being notably much more abundant (16,20). Type I 21U-RNAs in *C. elegans* are encoded by over 15,000 individual mini-genes organized within two large clusters at Chromosome IV (15-18). Promoters of many type I piRNA genes contain an 8-nt consensus Ruby motif which is associated with chromatin factors including PRDE-1, SNPC-4, TOFU-4 and TOFU-5 (21-24). Type II piRNA genes share transcription start sites with actively transcribed genes across the genome but lack the Ruby motif (16). Unlike other organisms, piRNA precursors in *C. elegans*, known as capped small RNAs (csRNAs), are relatively short, typically spanning 25-29 nts in length (16). Akin to other species, *C. elegans* piRNA biogenesis involves nucleolytic processing at both the 5’ and 3’ termini (1-3). The 5’ cap and the first 2 nts of csRNAs are removed by a trimeric Schlafen-domain nuclease (25). A few nucleotides are resected from their 3’ ends, followed by 2’–*O*–methylation catalyzed by the methyltransferase HENN-1 (26-29).

Despite the variations in lengths and sequences across species, piRNAs in worms, insects, and mammals all undergo 3’ trimming. For instance, our previous investigations led to the identification of PARN-1 as the specific exonuclease responsible for generating the mature 3’ ends of piRNAs in *C. elegans* (28). In *parn-1* mutant worms, piRNAs with 3′ extensions accumulate, while their 5’ cap and first 2 nts are correctly removed (28). A parallel study showed that Poly(A)-specific Ribonuclease-Like Domain Containing 1 (PNLDC1), a member of the PARN family, trims piRNA in *Bombyx mori* (30). Subsequent studies demonstrated that PNLDC1 acts on piRNAs in mice (31-33). Notably, exome sequencing of infertile male patients identified mutations in PNLDC1, linking these mutations to defects in piRNA processing and azoospermia (34,35). Collectively, these studies emphasize the importance of piRNA 3’ nucleolytic processing across various species.

PRG-1/piRNA complexes target a broad spectrum of germline transcripts as well as certain transposable elements (10,36-38). The targeting by PRG-1/piRNAs sets off a cascade of events. RNA-dependent RNA Polymerases (RdRPs) are recruited to target RNAs which serve as templates for the production of secondary small interfering RNAs (siRNAs). These siRNAs are typically 22 nts long and bear a 5’ guanine residue, hence referred to as 22G-RNAs (39-41). 22G-RNAs are then loaded onto a large group of worm-specific Argonautes (WAGOs) (10,11,36-38,42). Similarly, exogenous double-stranded RNAs (dsRNAs) can also induce the production of WAGO 22G-RNAs. In this process, endonuclease RDE-8 plays a pivotal role in cleaving the target RNAs (43). RDE-3, a terminal nucleotidyltransferase (TENT) (44), adds alternating uridine (U) and guanosine (G) nucleotides to the 3’ termini of cleaved transcripts, forming what is known as polyUG tails (45,46). RdRPs are subsequently recruited to the RNAs bearing polyUG tails, utilizing them as templates to synthesize 22G-RNAs (45). This amplification cycle serves as a potent mechanism for propagating and maintaining epigenetic silencing across generations (45,47). Thus far, RDE-3 and RdRPs are believed to act on long RNAs that are targeted by piRNAs or dsRNAs.

Here we report that *C. elegans* deficient for PARN-1 leads to the accumulation of a unique class of RNAs that are antisense to piRNAs, referred to as anti-piRNAs. The formation of anti-piRNAs involves a series of enzymatic activities. RDE-3 catalyzes the addition of non-templated UG dinucleotide and polyUG sequences at the 3’ termini of untrimmed piRNAs, though the UG additions are not a prerequisite for the production of anti-piRNAs. EGO-1, one of the functional RdRPs in *C. elegans*, utilizes untrimmed piRNAs as templates for synthesizing anti-piRNAs. Furthermore, a seed-gate structure within PRG-1 appears to limit the elongation of anti-piRNAs, resulting in 17-19 nt anti-piRNAs. In conclusion, our findings provide mechanistic insight to enzymatic activities of RDE-3 and EGO-1 and highlight the indispensability of piRNA 3’ processing.

## MATERIALS AND METHODS

### Caenorhabditis elegans strains

The Bristol strain N2 was designated as the wild-type *C. elegans* strain. Other strains used in this study can be found in Supplementary Table S1. All strains were grown on Nematode Growth Media (NGM) supplemented with OP50 or RNAi *E. coli* food and grown at 20°C unless otherwise indicated.

### CRISPR/Cas9 genome editing

PRG-1^SG_mut^ strains were generated using CRISPR/Cas9 genome editing and single stranded DNA donors in the *parn-1* mutant background (48). Cas9 loaded with gRNA targeting the PRG-1 seed gate genomic sequence was microinjected to *parn-1* adult animals along with ssDNA donors with 35 homology sequences. The PRF4 vector containing the dominant allele of *rol-6* served as an injection marker (48).

### RNA interference by feeding of double stranded RNA

The HT115 RNAi feeding strains were picked from the *C. elegans* Ahringer RNAi library (49). For RNA purification experiment 60,000 synchronized L1 larvae were platted on 150 mm NGM plates containing 50 μg/ml ampicillin and 5 mM IPTG seeded with HT115 bacteria against target genes. After reaching adulthood animals were collected with M9 and subjected to RNA extraction and small RNA enrichment.

### PRG-1 immunoprecipitation and RNA ligation

60,000 synchronous N2 and *parn-1* adult were collected, suspended and homogenized in one volume of IP buffer (110 mM Potassium acetate, 2 mM magnesium acetate, 0.1% Tween 20, 0.5% Triton, 1mM DTT, 20 mM HEPES-KOH, pH 7.5) with complete protease inhibitors (Roche). Cell extracts were cleared by two rounds of centrifugation at 14,000 x g for 10 min. Cell extracts were incubated with 12 μg anti-PRG-1 antibody (15) and 60 μl Protein A/G dynabeads (Thermo Fisher Scientific) at 4°C for 3 h. Beads were washed three times with IP buffer with protease inhibitors and once with wash buffer (150 mM NaCl, 2 mM magnesium acetate, 50 mM Tris-HCl pH 7.5). Beads were incubated with Tobacco Acid Pyrophosphatase (Epicentre) and washed once with wash buffer. In a 150 μl reaction, beads were incubated with 0.5 U/ μl T4 RNA ligase 1 (NEB) in 1x ligase buffer (containing 1 mM ATP, 1 U/uL RNase inhibitor and 10% PEG-8000) on the rocker at 16°C overnight. RNA was recovered by treatment with proteinase K (2.0 mg ml^-1^ in 0.5% (w/v) SDS, 40 mM EDTA, 20 mM Tris-HCl, pH 7.5) at 50°C for 10 min, followed by extraction with TRI Reagent (Sigma) and ethanol precipitation. To enrich the ligated product, RNAs ranging from 26 to 50 nts were purified from the 15% polyacrylamide/7M urea gel and subject to small RNA cloning.

### RNA extraction and small RNA enrichment

60,000 L1 larvae were plated to 150 mm NGM plates supplemented with 2 mL of concentrated OP50 food. Approximately 64 hours after plating L1 larvae, day 1 adults with eggs were collected from plates using M9 solution. Harvested animals were washed twice with M9 and once with water. Animals were then suspended in TRI Reagent solution (Sigma) and frozen at -80°C until RNA extraction was performed. To extract RNAs animals suspended in TRI Reagent solution were thawed and lysed using a Bead Mill (Thermo Fisher Scientific). Bromo-chloropropane (Sigma) was added to the lysed worms and centrifuged at 12,000 x g for 15 minutes. Following centrifugation, the aqueous phase was pipetted to an equal volume of cold isopropanol and placed at -20°C for 1 hour to precipitate RNAs. RNAs were pelleted by centrifugation at 20,000 x g for 15 minutes at 4°C. Total RNAs were washed with 75% EtOH twice and suspended in RNAse free water. Small RNAs were enriched from total RNAs using the MirVana miRNA isolation kit according (Thermo Fisher Scientific).

### Oxidation of small RNAs with NaIO_4_

5 μg small RNAs were oxidized with 25 mM sodium periodate (Fisher Scientific) in borax/boric acid buffer (0.06 M borax, 0.06 M boric acid, pH 8.6) in the dark at 20°C for 30 minutes. 20 μL of 50% glycerol was added and incubated for an additional 15 minutes to quench the sodium periodate. Oxidized RNAs were eluted from the reaction using EtOH precipitation at -20°C. Eluted RNAs were suspended in RNAse free water and subsequently used in small RNA cloning.

### Small RNA-sequencing library preparation

Small RNAs were treated with RNA phosphatase PIR-1 to remove γ and β phosphates from the 5′ triphosphorylated RNAs (50). Monophosphorylated RNAs were ligated to 3′ adapters ( rAppAGATCGGAAGAGCACACGTCTGAACTCCAGTCA/3ddC/, IDT) using T4 RNA ligase 2 in 25% PEG 8000 (NEB) at 15°C overnight. A 5′ adapter (rArCrArCrUrCrUrUrUrCrCrCrUrArCrArCrGrArCrGrCrUrCrUrUrCrCrGrArUrCrU, IDT) was then ligated to RNAs to the product using T4 RNA ligase 1 (NEB) for 4 hours at 15°C. Ligated products were reverse transcribed using SuperScript IV Reverse Transcriptase (Thermo Fisher Scientific) to convert RNA to cDNA libraries. cDNA libraries were amplified by PCR and subsequently sequenced on an Illumina Novaseq platform (SP2 x 50 bp) at the OSU Comprehensive Cancer Center genomics core.

### Small RNA-sequencing data analysis

Adapters and reads with poor quality were removed using TrimGalore (51). Identical sequences were counted and collapsed into fasta format using a custom shell script. Processed reads were aligned to a reference containing rRNAs, tRNAs, snRNA and snoRNAs. Reads aligning to this reference were excluded from downstream analysis as they likely represent contaminating RNA degradation products and can skew sample normalization. Reads were then aligned to the genome reference (WormBase, WS279) as well as a reference containing the sequences of exon-exon junctions using Bowtie (52). Reads aligning to the genome or junction reference were converted to bed format using BEDOPS (53). Alignments were intersected to a bed file containing gene annotations using BEDtools intersect (54). Reads overlapping with genomic features were filtered according to the following rules using a custom python script: 1) 22G/A siRNAs: reads mapping antisense to protein coding gene, lincRNA, pseudogenes or transposable element exons with a length of 21-23 nt and containing a 5’ G or A. The read must align perfectly but may map to up to 1000 genomic locations. 2) piRNAs: reads mapping sense to piRNA genes, containing a 5’ U of any length. The 5′ end of the read must map within 0,1, or 2 nucleotides relative to the annotated 5′ end of the piRNA gene. Reads mapping to piRNAs must map perfectly and uniquely. 3) Anti-piRNAs: reads mapping antisense to piRNAs. No filters were placed on 5′ nt identity or length. The read must align perfectly and uniquely. Anti-piRNAs were further filtered after preliminary analysis to be 17-19 nucleotides in length with a 5′ G/A. 4) miRNAs: Reads mapping to annotated miRNAs in the sense orientation with any 5′ nt and length. Following this filtration raw read counts were normalized to the total number of reads mapping to the genome or miRNAs, and then scaled to 1,000,000 total reads. The per-gene read count was then aggregated for each gene in each sample and used in downstream analysis. To generate BigWig files, bedfiles containing normalized reads passing filters for each genomic feature described above were converted to BedGraph files using BEDOPS (53). BedGraph files were converted to BigWig files using tools from UCSC (55). BigWig files were visualized in IGV (56).

### Analysis of PRG-1 IP and RNA ligation data

Raw reads from PRG-1 IP/RNA ligation experiments were processed as described above with unaligned reads retained for downstream analysis. Retained reads were aligned to a piRNA reference in which the annotated 3′ end of piRNAs was extended 50 nt. Alignments were filtered to select only those that: 1) mapped in the sense orientation, 2) mapped within position 0, 1, or 2 relative to the annotated 5′ terminus of the reference and 3) contained a tail or unaligned sequences that were at least 10 nt in length. Wild-type piRNA sequences had to precisely span 21 nucleotides, while *parn-1* samples allowed sense sequences of 21 nucleotides and longer. Unaligned sequences were subjected to an additional re-alignment step, demanding an antisense orientation to the piRNA reference. The identification of genuine piRNA::anti-piRNA ligation products was achieved through a custom Python script, merging reads containing both sense and antisense piRNA mapping sequences.

### Analysis of non-templated nucleotide additions

Processed and collapsed fasta reads were aligned to the genome reference (WormBase, WS279) allowing zero mismatches. Reads failing to align underwent a secondary alignment with Tailor (57), followed by conversion to bed files and assignment to specific genomic coordinates using BEDtools intersect (54). The same rules used for filtering reads overlapping genomic features described above were also used in the analysis of tailed reads. After annotating genes associated with tailed reads, further filtration involved: 1) retaining reads with 1-nucleotide tails. 2) for reads having tails of 2 to 5 nucleotides, requiring the edit distance (e.g. number of mismatches between the reference and read) to be equal to the length of tails. Reads failing this but being at least 3 nucleotides long were retained if their tail consisted of two alternating nucleotides (e.g., GUG/UGU). 3) preserving reads of at least 5 nucleotides entirely composed of a single nucleotide or two perfectly alternating nucleotides. Tailed reads meeting these criteria were normalized using the same constants as perfectly matched reads, as previously described. Tailed reads underwent further noise reduction by considering their presence in both sequencing replicates and the abundance of perfectly matched counterparts mapping to the corresponding genes. To filter according to abundance the Z score of perfectly matched reads mapping to a given gene was calculated in each genetic background, genes with a Z score ≥ 1 were retained for downstream analysis.

### CLASH data analysis

Published PRG-1 CLASH data were analyzed as previously described (58). Briefly, Adapters and reads with poor quality were removed using TrimGalore (51). Processed reads were collapsed to and aligned to the *C. elegans* reference. After identifying CLASH chimeras, the ΔG (kCal/mol) of piRNA::target interactions were calculated using tools from the Vienna RNA package (version 2.3.5) (59). To ensure the inclusion of confident piRNA::target interactions, those with a ΔG exceeding -15 kCal/mol were excluded from subsequent analysis. 1,650 piRNAs with anti-piRNA matching reads had detectable CLASH counts (58). Equitable comparisons were conducted by randomly sampling an equal number of piRNAs from the overall piRNA pool, ensuring that their abundance distributions closely matched those of the 1,650 piRNAs in our experimental group. This random sampling process was iterated 1,000 times to obtain the distribution of median CLASH count per piRNA for each sampling iteration, which was compared to the experimental set.

### Statistics and quantification

Analysis was preformed using published tools specified in the MATERIALS AND METHODS, custom R, Python, Nextflow and Shell scripts. Data were tested for normality before preforming parametric statistical tests. Statistical tests used are indicated in the Figure Legends.

## RESULTS

### anti-piRNAs accumulate in *parn-1* mutants

Acting as a 3’ to 5’ exonuclease, PARN-1 generates the mature 3’ ends of piRNAs in *C. elegans* (28). Worms deficient for PARN-1 accumulate untrimmed piRNAs with 3’ extensions and non-templated nucleotides at their 3’ ends (28,60). It is well-established that single-stranded precursors are produced from piRNA genes (1-4). In our previous analysis of small RNA sequencing (smRNA-seq) data, we exclusively focused on sense reads that mapped to piRNA-producing loci (28,60). By examining both sense and antisense reads that are uniquely mapped to the reference genome (Wormbase release WS279), we serendipitously discovered a previously unrecognized class of RNA species antisense to piRNAs in *parn-1* mutants, which we termed anti-piRNAs (Figure 1A). In the wild-type strain, small RNA reads predominantly matched the sense strand of piRNA genes (Figure 1A and 1B and Supplementary Table S2). In contrast, antisense reads were readily detected in *parn-1* mutants, although their abundance (reads per million total mapped reads or RPM = 541) was much lower than that of sense reads (RPM = 28,567) (Figure 1B). Both piRNA and anti-piRNA levels were dramatically reduced in *prg-1*; *parn-1* double mutants (Figure 1A and 1B and Supplementary Table S2), suggesting the accumulation of piRNA and anti-piRNAs depends on PRG-1 protein.

**Figure 1.**
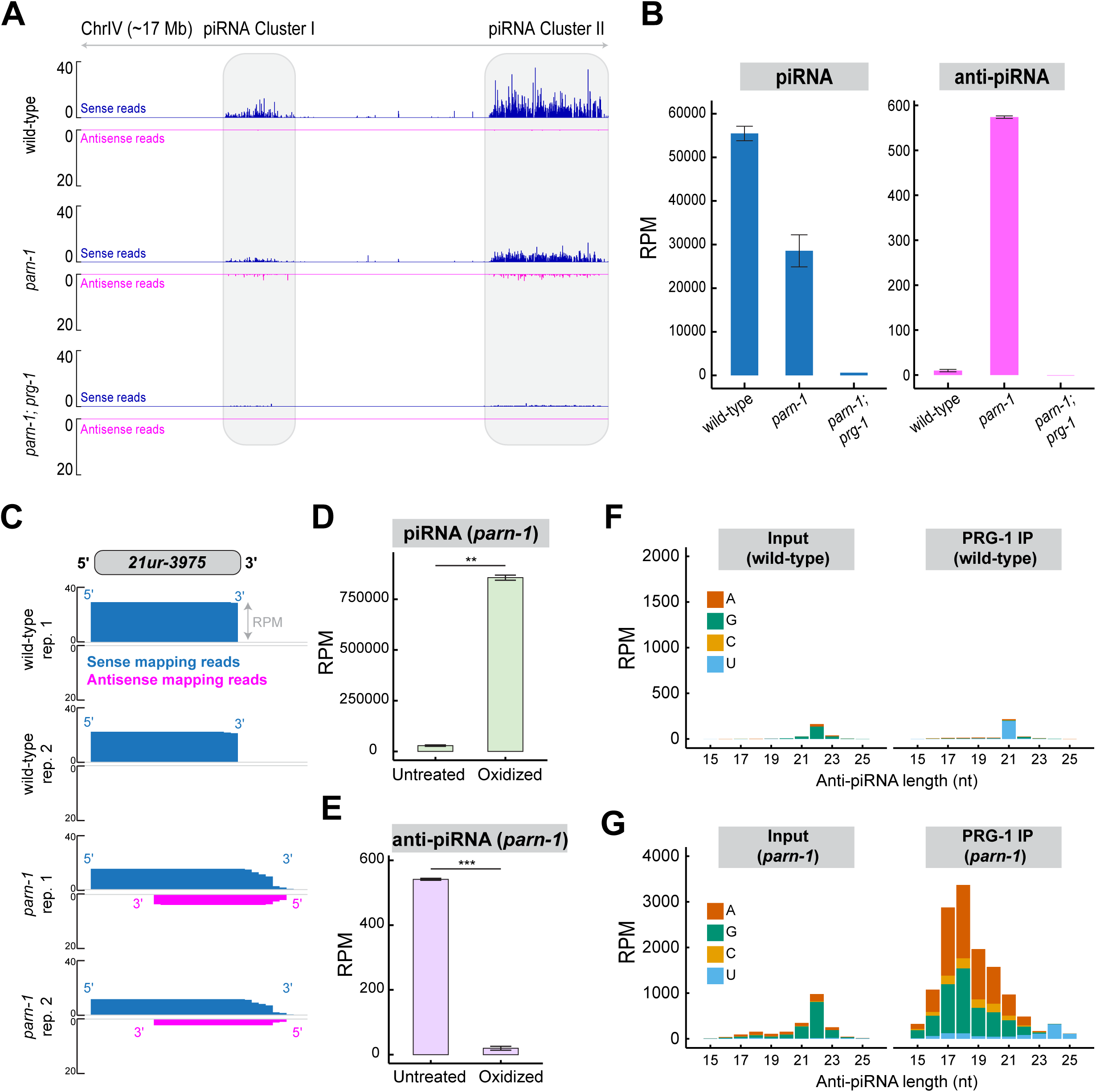
anti-piRNAs are produced in *parn-1* mutants. A) Browser view showing reads mapping sense and antisense to piRNA loci in wild-type, *parn-1* and *parn-1; prg-1* mutants. Sense sequences are shown in blue, antisense sequences are shown in magenta. *C. elegans* piRNA clusters are highlighted with grey boxes. Data are presented as the mean of (n = 2) biological replicates for wild-type and *parn-1*. And *parn-1; prg-1* data come from a single sample. Note that scales for sense and antisense reads are adjusted differently. B) Barplots showing the total level of piRNA (blue), and anti-piRNA matching reads (magenta) in wild-type, *parn-1* and *parn-1; prg-1* mutants. Data are displayed as the mean ± standard deviation of 2 biological replicates for wild-type and *parn-1*. *parn-1; prg-1* is comprised of a single biological replicate. C) Browser view showing sense and anti-piRNA reads mapping to 21UR-3975. In wild-type and *parn-1* mutants. Data are displayed as independent biological replicates. D) Barplots showing piRNA level under untreated and oxidizing conditions in *parn-1* mutants. Data are displayed as the mean ± standard deviation of 2 biological replicates. p < 0.01 **, two-sided T-test. E) Same as (D) but showing anti-piRNA level. F) Barplots showing the RPM of length and first nucleotide of antisense sequences in wild-type input and PRG-1 IP samples. Data are obtained from one biological replicate. G) Same as (F) but in *parn-1* mutants.

SmRNA-seq from *parn-1* mutants revealed that anti-piRNAs originate from 1,934 piRNA loci (Supplementary Table S3). We next inspected specific examples. Consistent with previous findings (28,60), piRNAs such as 21UR-3975 were primarily 21 nts in wild-type while untrimmed 21UR-3975 with 3’ extensions are found in *parn-1* mutants (Figure 1C). Anti-piRNAs were exclusive to *parn-1* mutants and were absent in wild-type strains (Figure 1C). Curiously, 5’ ends of anti-piRNAs initiated from various positions, while their 3’ ends mapped almost exclusively within the annotated piRNA gene (Figure 1C). In both wild-type and *parn-1* animals, piRNAs are known to bear 2’–*O*–methylation at their 3’ termini (26-29,60). To determine the methylation status of anti-piRNAs, we performed sodium periodate-mediated oxidation assays and sequenced oxidation-resistant RNA species. RNAs with terminal 2’-O-methylation are resistant to oxidation, while unmethylated RNAs possess vicinal diols at their 3’ termini, rendering them susceptible to oxidation and poor substrates for small RNA cloning. In *parn-1* mutants piRNAs were enriched by oxidation upon normalization to total number of reads (Figure 1D), suggesting untrimmed piRNAs possess terminal 2’-O-methylation (28,60). However, anti-piRNAs were strongly depleted by oxidation (Figure 1E), suggesting the absence of terminal 2’-O-methylation at anti-piRNAs.

The observation that the accumulation of anti-piRNAs requires PRG-1 prompted us to investigate whether anti-piRNAs are loaded onto PRG-1 (Figure 1A and 1B). To this end, we recovered PRG-1 complexes by immunoprecipitation (IP) from wild-type and *parn-1* extracts and subsequently sequenced PRG-1-bound small RNAs. In line with previous findings (28), wild-type piRNAs displayed characteristic 5’ U and a length of 21 nts, while untrimmed piRNAs with 5’ U were enriched in *parn-1* mutants from PRG-1 IP (Supplementary Figure S1A and S1B).

Next, we assessed antisense reads mapping to piRNA-producing loci. Some basal levels of 22G-RNA reads were detected in the input of the wild-type sample, but these were absent in the PRG-1 IP (Figure 1F and Supplementary Figure S1C). Additionally, we observed 21U-RNA reads that overlapped with piRNA-producing loci in the wild-type PRG-1 IP (Figure 1F and Supplementary Figure S1D). For instance, 21U-RNA reads mapped to the reverse strand of *21ur-11958* (Supplementary Figure S1D). However, the characteristic Ruby motif was found in the approximately 40-nt upstream of both 21U-RNA sequence and *21ur-11985* (Supplementary Figure S1D). We conclude these 21U-RNA reads are likely originated from unannotated piRNA genes in the complementary strand. These unannotated piRNA genes were also reported by a recent study (61). In stark contrast to the wild-type strain, PRG-1 IP from *parn-1* mutants revealed a substantial enrichment of anti-piRNAs (Figure 1G). Notably, these anti-piRNAs predominantly initiated with A or G residues (Figure 1G). Moreover, the size distribution of anti-piRNAs exhibited a broad range, peaking at 17-19 nucleotides (Figure 1G). We conclude that the size distribution and the 5’ nucleotide composition of anti-piRNAs are distinct from the well-characterized 21U-RNAs or 22G-RNAs.

### piRNA::anti-piRNA duplexes are loaded onto PRG-1

We proceeded to develop an approach to probe the formation of piRNA::anti-piRNA duplexes within PRG-1 by combining PRG-1 IP and RNA ligation (Figure 2A). Briefly, anti-PRG-1 antibody was used to enrich the PRG-1 complex (15). The purified complex was treated with Tobacco Acid Pyrophosphatase (TAP), an enzyme converting 5’ capped or polyphosphorylated RNA into 5’ monophosphorylated RNA. Proximal RNAs within the PRG-1 complex were then ligated to create hybrids. To enrich these hybrids, we isolated RNAs longer than 26 nucleotides, which were subsequently subjected to small RNA cloning and deep sequencing (Figure 2A).

**Figure 2.**
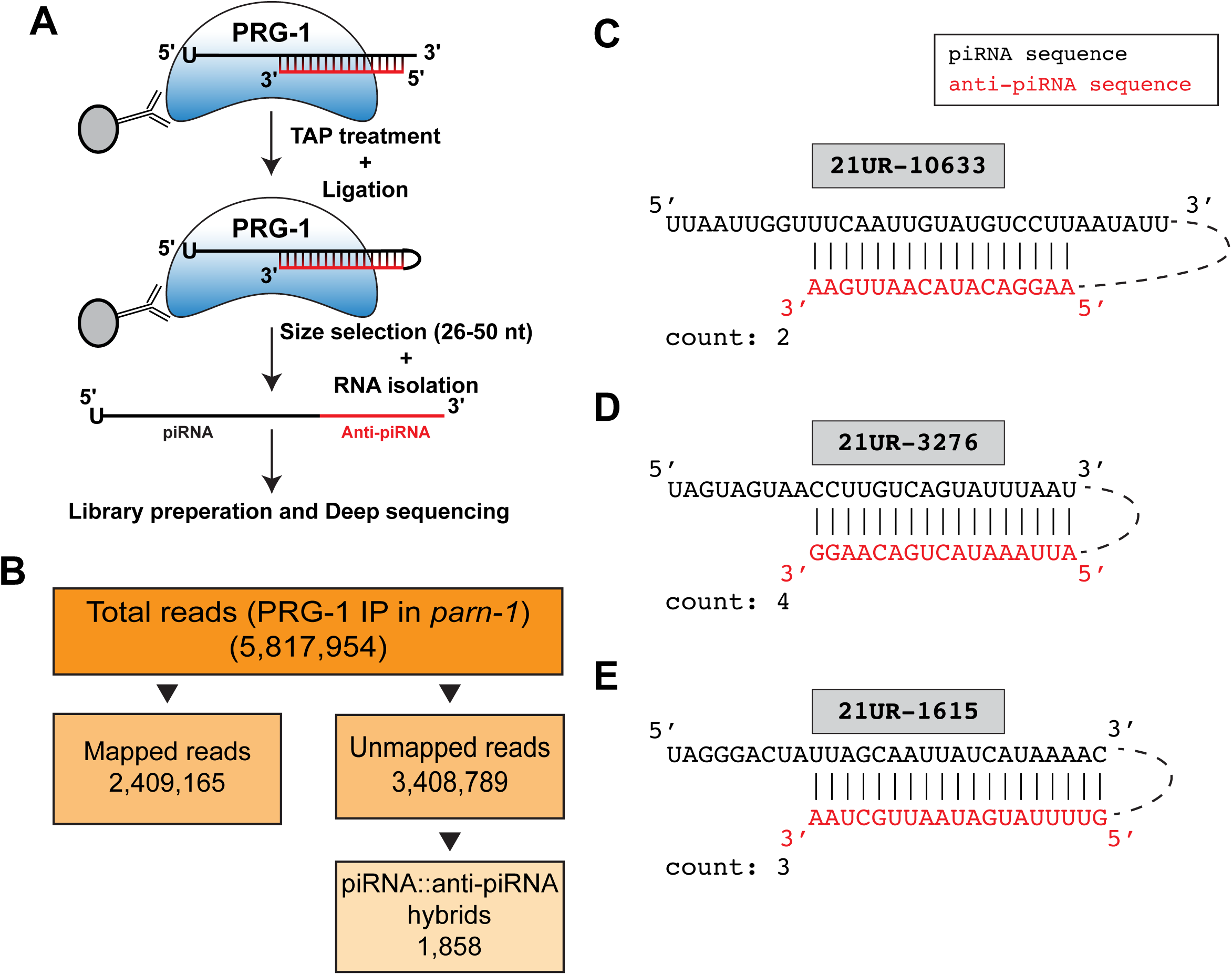
piRNA::anti-piRNA duplexes are loaded onto PRG-1. A) Schematic illustrating PRG-1 IP followed by ligation. Following PRG-1 IP RNAs bound to PRG-1 were treated with TAP. Adjacent RNAs were ligated together using RNA ligase. Chimeric RNAs were then subjected to RNA isolation, adapter ligation and deep sequencing. B) Flow chart showing library statistics and the number of piRNA::anti-piRNA chimeras from PRG-1 IP/ligation in *parn-1* mutants. C) Specific example showing sequencing reads of ligation between 21UR-10633 and an associated anti-piRNA (red). The raw sequence count of this chimera measured by smRNA sequencing is shown. D) Same as in (C) but showing locus 21UR-3276. E) Same as in (D) but showing locus 21UR-1615.

Our experimental effort in wild-type and *parn-1* backgrounds yielded ∼2.77 million and ∼5.82 million sequence reads, respectively (Figure 2B and Supplementary Figure S2). Of these reads, a substantial portion mapped to the genome reference (Figure 2B and Supplementary Figure S2). Among reads that failed to map to the genome, we searched for hybrid reads composed of a piRNA sequence and anti-piRNA sequence. No piRNA::anti-piRNA hybrids were identified in wild-type (Supplementary Figure S2). In *parn-1* mutants, we recovered 1,858 hybrid reads, which were distributed across 1,533 piRNA-producing loci (Figure 2B). In silico folding on hybrid reads revealed the evidence for the presence of piRNA::anti-piRNA duplexes (Figure 2C-E). In summary, our smRNA-seq data and bioinformatic analyses uncovered anti-piRNA species that are generated upon the loss of PARN-1. Anti-piRNAs are 17-19 nts in length, forming duplexes with untrimmed piRNAs, and loaded onto PRG-1. Furthermore, we showed that anti-piRNAs possess 5’ A or G, but do not contain 2’–*O*–methylation 3′ residues, distinguishing them from their piRNA counterparts.

### RDE-3 modifies 3’ termini of untrimmed piRNAs

Next, we aimed to identify the enzymatic factors responsible for anti-piRNA production. We hypothesized that RNA polymerases may play a role in generating anti-piRNAs, given their preference for initiating transcription with a purine (A or G) as the first base in nascent transcripts (62-64). Initially, we considered the possibility that anti-piRNAs, akin to piRNA precursors, are produced by RNA polymerase II. However, by mining transcription start site databases (16,65), we found anti-piRNA reads did not coincide with transcription start sites. This led us to explore an alternative hypothesis: anti-piRNAs are synthesized by RNA-dependent RNA polymerases (RdRPs) using piRNAs as templates. It is known that exogenous dsRNAs or endogenous triggers can induce the cleavage of target RNAs (43,45). Following cleavage, RDE-3 adds polyUG tails to the 3’ termini of cleaved RNAs (45,46) (Figure 3A). RdRPs, including RRF-1 and/or EGO-1, initiate the synthesis of 22G-RNAs, which are templated directly from polyUG RNAs (45) (Figure 3A). A speculative but intriguing possibility is that the 3′ termini of untrimmed piRNA could not be properly accommodated by PRG-1 and therefore are susceptible to RDE-3 and RdRP(s).

**Figure 3.**
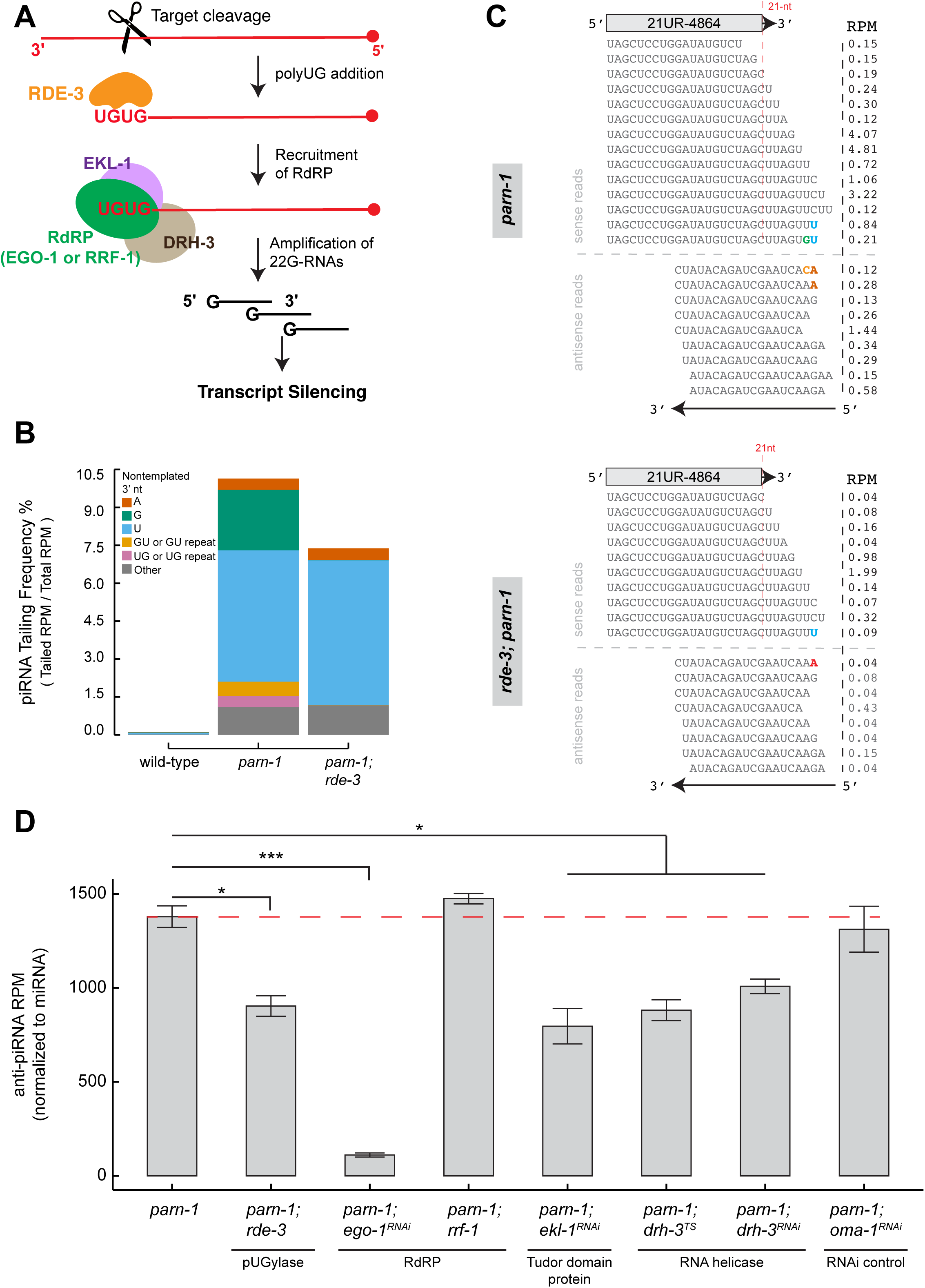
anti-piRNAs are targeted by RDE-3 and EGO-1. A) Schematic illustrating 22G-RNA synthesis in wild-type *C. elegans*. Following target cleavage RDE-3 is recruited to the 3’ end of target RNAs and adds stretches of polyUG repeats These repeats recruits the RdRP complex composed of EGO-1 or RRF-1 (RNA-dependent RNA polymerase), EKL-1 (Tudor domain protein), and DRH-3 (RNA helicase) for 22G-RNA production. B) Barplot showing the frequency of non-templated nucleotide additions (3’ tailing) to piRNAs in wild-type, *parn-1* and *parn-1; rde-3* mutants. piRNA tailing frequency is derived using the following equation: 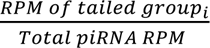 where *RPM tailed group_i_* is the abundance of reads with tail *i* (mono-A, mono-G, mono-C, mono-U, GU or GU repeats, UG or UG repeats, and others), and *Total piRNA RPM* is the total abundance of all reads mapping to piRNA loci (perfectly matching and tailed reads). Data are displayed as the mean of 2 biological replicates. C) Specific example showing the sequence of sense piRNA and anti-piRNA sequences mapping to 21UR-4864. The graphic shows both perfectly matched and tailed piRNAs, anti-piRNAs, as well as anti-piRNAs mapping to tailed sense piRNAs. Non-templated nucleotides are shown in blue (U), red (A), orange (C) and green (G). Nucleotides matching the endogenous sequence are shown in grey. Anti-piRNAs with nucleotides mapping to tailed sense piRNAs are highlighted with a grey bar for clarity. RPMs are displayed as the mean of 2 biological replicates. D) Barplot showing the level of anti-piRNAs (normalized to miRNA) in *parn-1, parn-1; rde-3, parn-1; ego-1^RNAi^, parn-1; rrf-1, parn-1; ekl-1^RNAi^, parn1; drh-3^TS^, parn-1; drh-3^RNAi^* and *parn-1; oma-1^RNAi^.* Data are displayed as the mean ± standard deviation of 2 biological replicates. p < 0.05 *, p < 0.001 ***, two-sided T-test.

Our previous study revealed abundant 3’ non-templated nucleotide additions, also known as 3’ tailing, in *parn-1* mutants (60). At that time, our customized bioinformatic pipeline allowed us to detect up to three occurrences of non-templated nucleotides (60). For this investigation, we employed the short-read aligner Tailor, which allowed us to unbiasedly determine non-templated nucleotide additions to RNAs regardless of their lengths (57). In line with prior findings (60), a small fraction of piRNAs exhibited 3’ non-templated additions in wild-type worms. In *parn-1* mutants, the frequency of non-templated nucleotide additions increased to 10.14% (Figure 3B). Mono-uridylation (mono-U) was the most abundant modification in both wild-type and *parn-1* strains, while mono-guanylation (mono-G) of piRNAs became abundant in *parn-1* mutants (Figure 3B) (60). Both UG dinucleotide/repeat (0.43%) and GU dinucleotide/repeat (0.58%) were detected in piRNAs upon loss of PARN-1 (Figure 3B). Strikingly, mono-G, UG, and GU additions were significantly reduced in *rde-3*; *parn-1* double mutants compared to their levels in *parn-1* single mutants, while mono-A and mono-U levels remained comparable between these two strains (Figure 3B).

Loss of mono-G in *rde-3*; *parn-1* mutants was unexpected, given that a previous cellulo-tethering assay suggested that RDE-3 catalyzes polyUG additions (46). To further investigate this matter, we assessed the nucleotide composition at positions one or two nucleotides upstream (-1 or -2 positions) of the 3’ non-templated nucleotides. (Supplementary Figure S3A and S3B). The Tailor pipeline defined the -1 or -2 nucleotides as templated nucleotides, as they matched the genome reference. However, they could also be non-templated nucleotides if the added 3’ tails matched the genome reference purely by chance. In the group of piRNAs with mono-A tails, there was an overrepresentation of A at both the -1 and -2 positions, implying that polyA may be added to the 3’ termini of untrimmed piRNAs (Supplementary Figure S3A and S3B). U was overrepresented at the -1 position in groups with mono-G and GU tails, but not in other groups (Supplementary Figure S3A). No such strong bias was observed at the -2 position (Supplementary Figure S3B). These findings suggest that U addition may precede G and GU addition. It is also possible that piRNAs terminating with U are preferred substrates for RDE-3.

Finally, we inspected specific piRNA-producing loci. For example, in *parn-1* mutants sense reads mapped to *21ur-4864* locus exhibited 3’ extensions and non-templated nucleotide additions including mono-U and GU. Antisense reads initiated from various positions. On some occasions they were complementary to sense reads with the non-templated nucleotides (Figure 3C). In agreement with the genome-wide analysis (Figure 3B), the non-templated GU addition at 21UR-4864 was absent in *rde-3*; *parn-1* mutants (Figure 3C). Taken together, these findings suggest that RDE-3 catalyzes 3’ non-templated UG dinucleotide and repeat additions to untrimmed piRNAs.

### EGO-1 uses untrimmed piRNAs as templates to initiate anti-piRNA production

To determine whether RdRPs are responsible for generating anti-piRNAs, we depleted individual RdRP components to assess their impact on anti-piRNA production. Two RdRPs, RRF-1 and EGO-1, function partially redundantly in the germ line to produce 22G-RNAs using endogenous transcripts or transposons as templates (39-41). RRF-1 and EGO-1 individually form a complex with the Tudor-domain protein EKL-1 and RNA helicase DRH-3 (40,66). While *rrf-1* is a nonessential gene, EGO-1, EKL-1 and DRH-3 are required for viability (39-41,66).

To deplete RdRP components in the *parn-1* mutant background, we utilized a loss-of-function allele of *rrf-1* (67), temperature-sensitive allele of *drh-3* (40), and performed knockdown of *ego-1*, *drh-3* and *ekl-1* using RNA interference (RNAi) (66). As expected, depletion of *rde-3*, *rrf-1 ego-1*, *drh-3* and *ekl-1* all resulted in a reduction in 22G-RNA levels (Supplementary Figure 3C). Since the degree of reduction varied across samples (Supplementary Figure 3C), we decided to normalize anti-piRNAs reads against microRNA reads instead of total mapped reads to accurately quantify anti-piRNA abundance (Figure 3D). While *rde-3*; *parn-1* mutants displayed the strongest depletion of 22G-RNAs (Supplementary Figure 3C), their anti-piRNA levels were only modestly decreased, suggesting RDE-3 contributes to anti-piRNA production but is not required for its production (Figure 3D). The deletion of *rrf-1* led to slight increase, albeit not statistically significant, in anti-piRNA levels. On the other hand, *ego-1* depleted animals exhibited ∼12.4-fold decrease in anti-piRNA levels relative to the control (Figure 3D), suggesting EGO-1 is the primary RdRP for anti-piRNA production. Consistent with the idea that EGO-1, EKL-1 and DRH-3 form a RdRP complex (40), depletion of either *ekl-1* or *drh-3* by RNAi resulted in the reduction of anti-piRNAs (Figure 3D). Taken together, our studies provide the evidence that the EGO-1-containing RdRP complex initiates the de novo synthesis of anti-piRNAs that are templated directly from untrimmed piRNAs.

### The seed-gate structure within PRG-1 hinders anti-piRNA elongation

Since the EGO-1/EKL-1/DRH-3 complex normally produces of 22G-RNAs (40), we wondered why it can generate anti-piRNAs that are 17 to 19 nts in length (Figure 1G). We first measured the distance between 5’ ends of piRNA (5’ U) and 5’ ends of anti-piRNAs (Figure 4A). While the piRNA size in *parn-1* mutants showed a broad distribution (22-29 nts) (28), 5’ piRNA to 5’ anti-piRNA distance exhibited a narrower range (25-29 nts) (Supplementary Figure S4A). This finding indicates that longer isoforms of untrimmed piRNA are preferred RdRP substrates. We then measured the distance between 5’ ends of piRNA and 3’ ends of anti-piRNAs (Figure 4A). The distance exhibited a narrow range and peaked at 9 nts (Figure 4B). In other words, the last nucleotide of anti-piRNAs primarily base-paired with the 10^th^ nucleotide of corresponding piRNAs (Figure 4B). This observation motivated us to test two models for the generation of anti-piRNA 3’ ends. 1) the RdRP complex uses entire piRNAs as templates to generate RNA duplexes and PRG-1 cleaves antisense RNAs at the phosphodiester bond linking nucleotides 9 and 10 (Figure 4C). 2) RdRP polymerization reaction is terminated due to the steric hindrance imposed by certain structure(s) within PRG-1 (Figure 4C).

**Figure 4.**
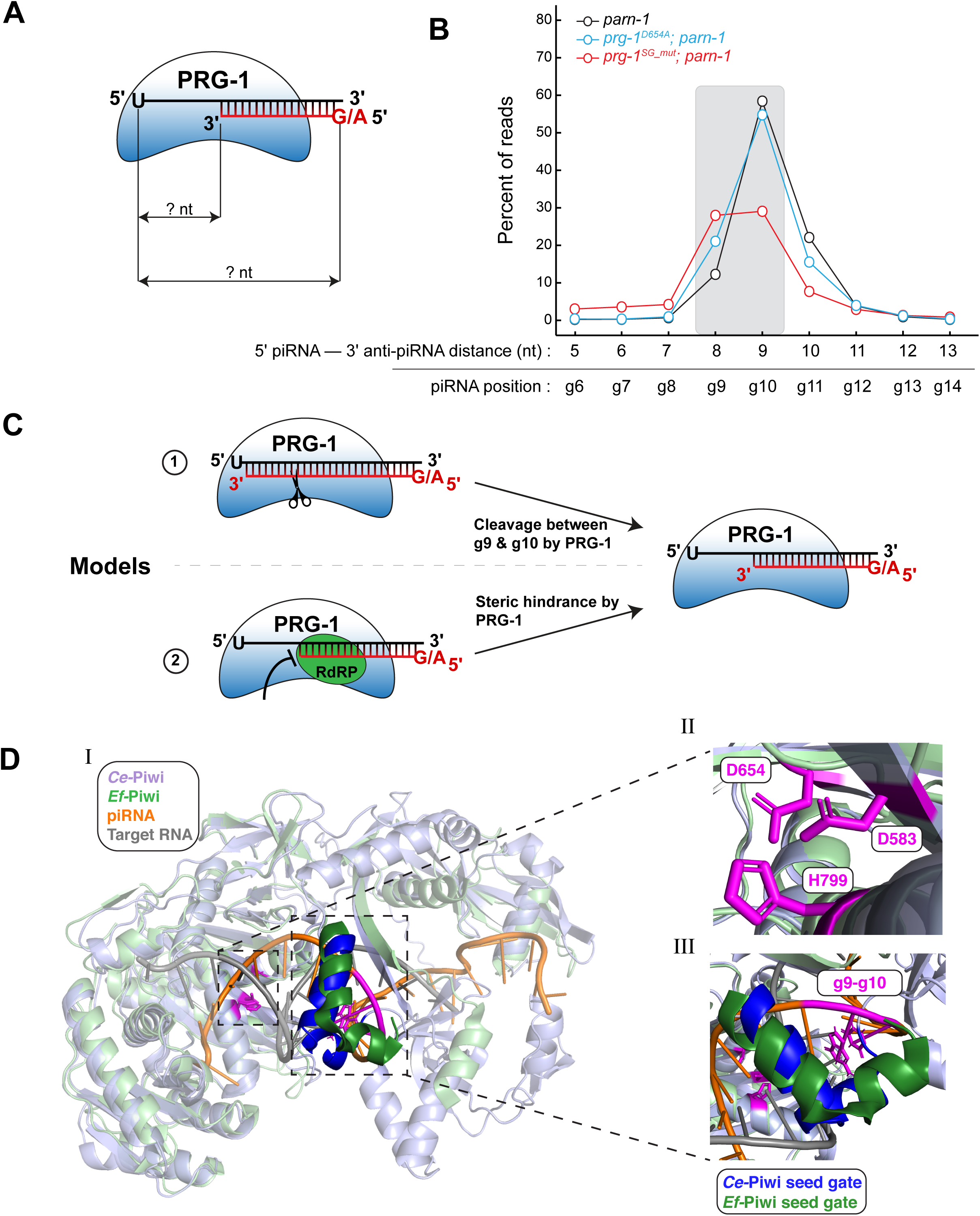
PRG-1 seed gate may block RdRP from synthesizing anti-piRNAs. A) Schematic illustrating the information derived from measuring the distance from the 5′ or 3′ ends of anti-piRNAs (red) to the 5′ ends of sense piRNAs (black). The 5′ to 3′ (piRNA to anti-piRNA) distance describes where anti-piRNA synthesis stops, conversely 5′ to 5′ distance describes where anti-piRNA synthesis begins. B) Line plot showing the 5′ to 3′ distance of piRNAs to anti-piRNAs in *parn-1, parn-1; prg-1^D654A^; parn-1; prg-1^SG_mut^*. Data are displayed as distances relative to the 5′ end of piRNAs (position 0). The y-axis is the percent of reads with the 5′ to 3′ distance on the x-axis. A grey box highlights a 5′ to 3′ distance of 8 and 9 nucleotides. Data are displayed as the mean of 2 biological replicates. g: guide RNA. C) Schematic illustrating two models by which 3′ ends of anti-piRNAs are generated. In the first model synthesis of anti-piRNAs (red) goes beyond g10 of piRNAs (black) and PRG-1 cleaves anti-piRNAs between g9 and g10. In the second model RdRP elongation is halted due to steric hinderance by PRG-1 protein itself. D) Cartoon representation of *C. elegans* PRG-1 (*Ce-*PIWI: Alphafold prediction) superimposed to *Ephydatia fluviatilis* (*Ef-*PIWI: Cryo-EM structure; PDB: 7KX9). Shown in orange and grey color are the piRNA and target RNA bound to *Ef-*PIWI, respectively (I). Zoomed-in cartoon showing the *Ce-*PIWI catalytic triad DDH, highlighted in magenta. In 4B D654 is mutated to alanine (II). Zoomed-in cartoon showing the *Ce-*PIWI seed gate (dark blue) and *Ef-*PIWI seed gate (dark green). Highlighted in magenta are g9-g10 of the piRNA. In 4B the PRG-1 seed gate (L332-T350) was replaced with a Gly_6_ linker (III).

To test the first model, we assessed anti-piRNA production in catalytically inactive PRG-1 mutants. PRG-1 contains evolutionarily conserved catalytic residues (DDH) essential for its slicer activity (42), although the slicer activity is dispensable for piRNA-mediated silencing (36,37,47). Previous in vitro assays have shown that PRG-1 slicer activity is abolished by mutating its DDH triad to DAH (37). CRISPR/Cas9 genome editing was used to introduce the DAH mutation (D654A) in the *prg-1* genomic locus in a previous study (Figure 4D) (47). We found that the anti-piRNA level in *prg-1* (D654A); *parn-1* was comparable to that in *parn-1* mutants (Supplementary Figure S4B). Importantly, the distance between 5’ ends of piRNA and 3’ ends of anti-piRNAs in *prg-1* (D654A); *parn-1* peaked at 9, similar to what was observed in *parn-1* mutants (Figure 4B). Collectively, these findings suggest that PRG-1 slicer activity does not play a role in the production of anti-piRNAs.

To explore the second model, we examined the PRG-1 structure that is predicted by AlphaFold (Figure 4D) (68). The AlphaFold program reported high confidence scores for the PAZ, MID, and PIWI domains of PRG-1. We superimposed the predicted PRG-1 structure to the cryo-EM structure of the Piwi-piRNA-target complex from the sponge Ephydatia fluviatilis (Ef-Piwi) (Figure 4D) (69). Within the Ef-Piwi, Drosophila Piwi, and silkworm Siwi structures, a distinct α-helix referred to as the seed-gate is present near the piRNA seed region (position 2 to 8) (69). This seed-gate has been observed to move upon binding to targets and is essential for targeting fidelity (69). The seed-gate structure was observed in PRG-1 (Figure 4D). Notably, it was in close proximity to the position 9 and 10 of piRNAs, where most of anti-piRNAs terminate (Figure 4D). We therefore tested the idea that the seed-gate structure might obstruct RdRP-mediated anti-piRNA elongation. Disruption of the seed-gate (PRG-1^SG_mut^ in which L332-T350 replaced with a Gly_6_ linker) led to a decrease in piRNA levels (Supplementary Figure S4B). Although it is important to note that this mutation could potentially alter the conformation of PRG-1, the disruption of PRG-1 seed-gate resulted in the production of longer anti-piRNAs and a one-nucleotide shift in the distance between the 5’ ends of piRNA and the 3’ ends of anti-piRNAs (Figure 4B). We conclude that the seed-gate within PRG-1 may impose steric hindrance that blocks RdRP activity, thereby hindering anti-piRNA elongation.

### piRNA-target interactions promote the generation of anti-piRNAs

It is noteworthy that only a subset of piRNAs (n = 1,934) in *parn-1* mutant strains were templated to generate anti-piRNAs (Supplementary Table S3). Furthermore, there was a poor correlation between the abundance of anti-piRNAs and their corresponding piRNAs (Pearson’s *ρ* = 0.05) (Figure 5A), suggesting the abundance of piRNA is not the determinant factor for anti-piRNA production. This raises the question of why certain piRNAs were selectively targeted by the RdRP complex. Previous studies demonstrated that base-pairing interactions between piRNAs and their targets can recruit RdRPs to target RNAs, which serve as templates to produce 22G-RNAs (10,11,36,37). This led us to explore the possibility that piRNA::target interactions might also facilitate the synthesis of anti-piRNAs by recruiting RdRPs to untrimmed piRNAs.

**Figure 5.**
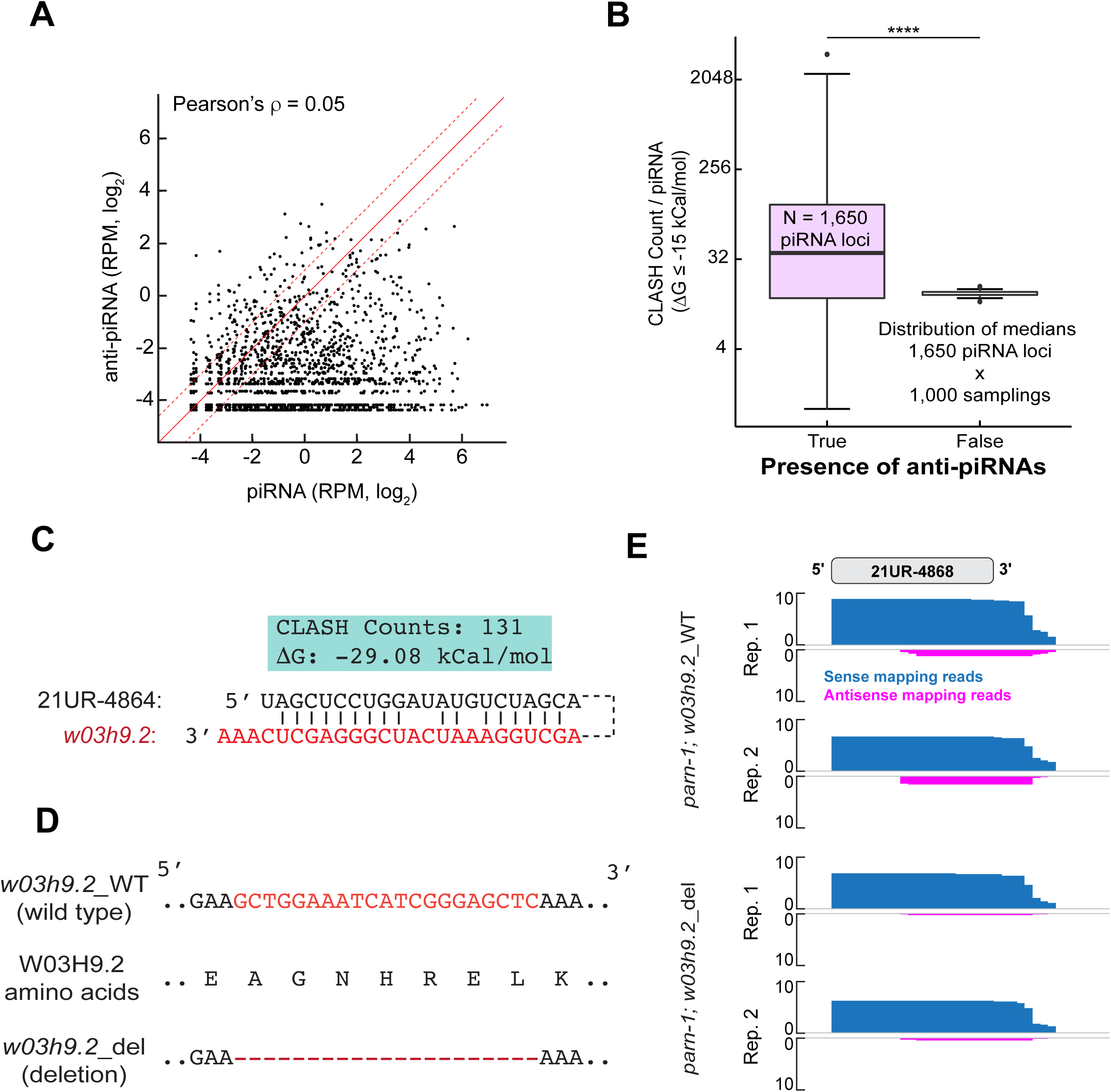
piRNA::target interactions facilitate anti-piRNA synthesis. A) Scatterplot comparing the level of sense and antisense sequences for a given piRNA (N = 1,934) in *parn-1* mutants. The solid red line represents no difference. The upper and lower dashed red lines represent a fold-difference of 2 and 0.5, respectively. Pearson’s correlation test was used to derive ρ. B) Boxplot showing the CLASH count per piRNA for piRNAs with anti-piRNA mapping sequences (N = 1,650) and the control set. The control set consists of random 1,650 piRNAs with comparable abundance to the experimental set but lacking anti-piRNA sequences. This random sampling process was repeated 1,000 times to obtain the distribution of median CLASH count. p < 0.0001 ****, two-sided T-test. C) Schematic illustrating the CLASH chimera between 21UR-4864 (black) and w03h9.2 (red). Shown in the blue box are the CLASH count detected for this interaction and the predicted free energy (1′G) of the interaction. D) Schematic illustrating the *w03h9.2* genomic locus, 21UR-4864 target site in w03h9.2 transcript (Red), the amino acids this sequence encodes, and deletion of the target site (*w03h9.2_del*) E) Genome browser view showing sense (blue) and anti-sense (magenta) reads mapping to 21UR-4864 in *parn-1*; *w03h9.2_WT* and *parn-1*; *w03h9.2_del*. Data are displayed as individual biological replicates.

The assessment of the target::piRNA interactome was performed using CLASH, an approach combining crosslinking, ligation of piRNAs and their targets, and subsequent sequencing of hybrid molecules (58,70). In silico folding was conducted to determine the base-pairing interaction within hybrid reads, and the Gibbs free energy (ΔG) of the most energetically favorable interaction can be subsequently deduced (Supplementary Figure S5A) (58). It is important to acknowledge that the CLASH experiment was performed in the wild-type background, but not in *parn-1* mutants (58). However, considering that the seed sequence of piRNAs largely governs piRNA::target interactions (58), we posited that hybrid reads derived from wild-type could serve as a proxy for piRNA::targets interactions in *parn-1* mutants. Therefore, we cross-referenced piRNAs with anti-piRNA sequences defined in this study with piRNA::target interactions previously revealed by the CLASH experiment (58). When focusing on robust piRNA::target interactions (ΔG ≤ -15 kCal/mol), we observed that out of the 1,934 piRNAs containing anti-piRNA sequences, 1,650 exhibited detectable CLASH counts. To create a control group for comparison, we randomly selected 1,650 piRNAs with comparable abundance to our experimental set but lacking anti-piRNA sequences. This random sampling process was repeated 1,000 times to obtain the distribution of median CLASH count (Figure 5B). Importantly, the CLASH counts for piRNAs with anti-piRNAs were found to be significantly higher than those in the control group (Figure 5B). This finding suggests a positive correlation between piRNA::target interactions and the synthesis of anti-piRNAs.

Next, we aimed at determining the causal relationship between piRNA::target interactions and anti-piRNA production. Our previous study established the base-pairing interaction between 21UR-4864 (piRNA) and endogenous *W03H9.2* transcript (target RNA) (60). This interaction was characterized by abundant CLASH reads and extensive base-pairing interactions (ΔG = -29.08 kCal/mol) (Figure 5C, D) (60). In this study, we discovered anti-piRNAs originating from the *21ur-4864* locus in *parn-1* mutants (Figure 5E). We reasoned if anti-piRNAs are indeed induced by base-pairing interactions between piRNAs and their targets, disrupting the interaction between 21UR-4864 and *W03H9.2* would negatively impact anti-piRNA production. Using CRISPR/Cas9 genome editing, we have generated a deletion allele of w*03h9.2* (*w03h9.2_*del) that specifically deleted the 21UR-4864 binding site from *W03H9.2* mRNA and removed seven amino acids from W03H9.2 protein (Figure 5D) (60). Consistent with the idea that piRNA induces 22G-RNA production (10,11,36,37), the deletion of 21UR-4864 target site (*w03h9.2_*del) resulted in a reduction of 22G-RNAs that are antisense to *W03H9.2* transcript in *parn-1* mutants (Supplementary Figure S5B). When examining the expression of anti-piRNAs that are antisense to 21UR-4864, we found that *w03h9.2_del* mutants displayed 2.95-fold reduction in anti-piRNAs levels compared to the control group (Figure 5E). Taken together, our findings provide evidence that piRNA::target interactions promote the production of anti-piRNAs in *parn-1* mutants.

## DISSCUSION

From nematodes to mammals, the maturation of piRNAs invariably involves 3’ nucleolytic processing (28,30-33). Our previous study demonstrated the evolutionarily conserved ribonuclease PARN-1 mediates piRNA 3’ trimming in *C. elegans* (28). In this study, we report that untrimmed piRNAs in *parn-1* mutants were modified by RDE-3 and templated by EGO-1/EKL-1/DRH-3 complex to produce anti-piRNAs (Figure 6). In wild-type, piRNA targeting may trigger the cleavage of target RNAs whose 3’ termini are subject to polyUG addition (45,46). RdRP complexes are recruited to PolyUG targets which serve as templates to produce 22G-RNAs that are subsequently loaded to WAGOs (Figure 6). In *parn-1* mutants, however, 3′ termini of untrimmed piRNA may fail to be accommodated properly by the PAZ domain of PRG-1. This results in the erroneous recognition and modification of these long untrimmed piRNAs by RDE-3. Upon recruitment, RdRP complexes not only use targets as templates to generate 22G-RNAs, but also use untrimmed piRNAs (with or without UG/GU additions) as templates to synthesize anti-piRNAs (Figure 6). Interestingly, while 22G-RNA production depends on RNA-dependent RNA polymerase EGO-1 and RRF-1, the synthesis of anti-piRNAs primarily relies on EGO-1 (Figure 3D). The elongation of anti-piRNAs is halted by the seed-gate structure of PRG-1, resulting in the synthesis of 17-19 nt anti-piRNAs that associate with the PRG-1 protein (Figure 6). The precise physiological significance of anti-piRNA production remains elusive. We speculate that the base-pairing interaction between anti-piRNAs and piRNAs can hinder piRNAs from recognizing their authentic targets. This could explain why untrimmed piRNAs in *parn-1* mutants are partially defective in 22G-RNA production and silencing of foreign sequences (28,60). Collectively, our findings underscore the critical role of nucleolytic processing at piRNA 3’ termini and unveils a novel class of small RNAs.

**Figure 6.**
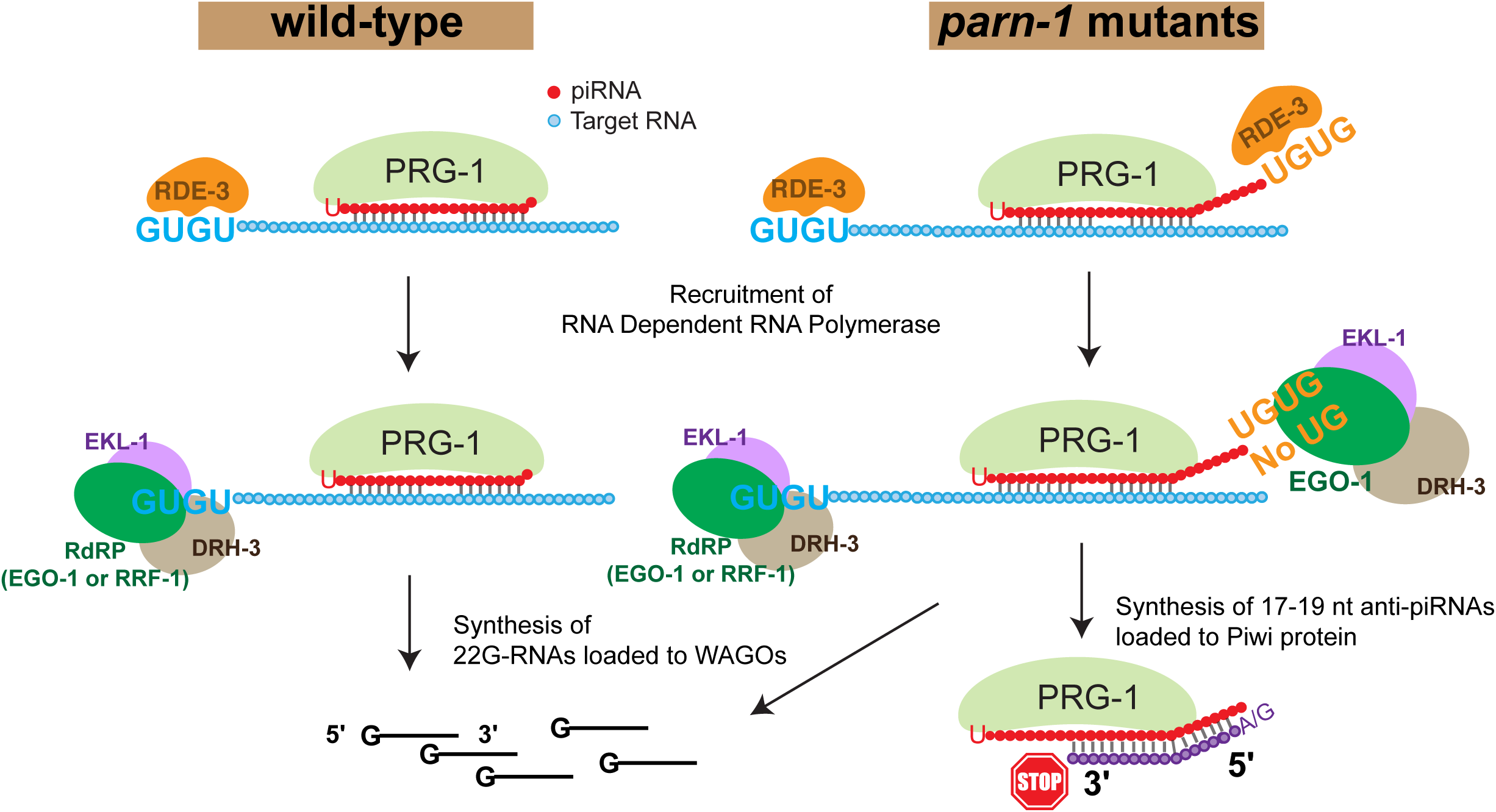
Model of anti-piRNA biogenesis. Schematic illustrating the synthesis of 22G-RNAs and anti-piRNA in wild-type and *parn-1* mutant strains.

Although anti-piRNAs were exclusively identified in the *parn-1* mutant background, their discovery sheds light on the enzymatic activities of RDE-3, RRF-1 and EGO-1. Previous studies described RDE-3 as the terminal nucleotidyltransferase adding alternating U and G to long RNAs during RNAi (45,46). Our data provide additional insights into RDE-3 substrates and activities. Firstly, RDE-3 can target small RNAs that are not properly processed, indicating that substrate lengths do not determine RDE-3 targeting. Secondly, RDE-3 has the capacity to catalyze UG dinucleotide additions (Figure 3B), which raises the possibility that RDE-3 may not exhibit high processivity in vivo. It is important to note that our small RNA cloning method may not capture piRNAs with long polyUG tails, as these long tails form atypical RNA quadruplex complexes which may hinder adaptor ligation (71). Alternatively, some cellular ribonucleases may shorten piRNA polyUG tails. To comprehensively characterize RDE-3’s enzymatic activities and define its substrate specificity, further investigations using biochemical assays are warranted.

Two RdRPs, RRF-1 and EGO-1, are thought to function partially redundantly in the production of 22G-RNAs (40). However, our findings suggest that they have a distinct role in anti-piRNA synthesis. Specially, RRF-1 is dispensable for anti-piRNA production, while EGO-1 emerges as the primary RdRP responsible for anti-RNA production (Figure 3D). Previous studies have showed that RRF-1 exhibited stronger binding to polyUG tails compared to EGO-1 (45,46). This suggests that polyUG tails may be crucial for recruiting RRF-1 to target RNAs but less so for EGO-1 recruitment (Figure 3D). This idea aligns with our observations of modest anti-piRNA depletion in *rde-3* mutants and robust anti-piRNA depletion in *ego-1* mutants. Furthermore, our data reveal that EGO-1 initiates anti-piRNA synthesis with either an A or G as the first base, occurring at a similar frequency (Figure 1G). When recruited to RNA targets, EGO-1 may generate secondary siRNAs starting with either A or G. The preference for 5’ G found in 22G-RNAs may be dictated by individual WAGO proteins. In support of this notion, a recent survey of *C. elegans* Argonaute proteins revealed that certain WAGOs (such as WAGO-1) predominantly associate with small RNAs initiating with G, while others (such as HRDE-1/WAGO-9) are associated with small RNAs initiating with A or G (61). In vitro biochemical experiments will be needed to establish the distinct activities of RRF-1 and EGO-1.

## AUTHOR CONTRIBUTION

B.P., and W.T. conceived and designed the experiments. B.P., W.T., and H.L.H., preformed the experiments. B.P., and W.T., analyzed the deep sequencing data. B.P., H.L.H., and W.T. wrote and edited the manuscript.

## ACKNOWLEDGEMENTS

We thank C. Mello for the support during the initiation phase of this project, S. Tu for assistance in initial data analysis, S. Kennedy for providing PRG-1 (D654A) mutants. The Ohio Supercomputer for supercomputing resources, the OSU Comprehensive Cancer Center genomics core for Illumina sequencing, Caenorhabditis Genetics Center for providing some of the *C. elegans* strains (P40OD010440), and the Ohio State University Center for the RNA Biology fellowship awarded to B.P.

## FIGURE LEGENDS

**Supplementary Figure S1.**
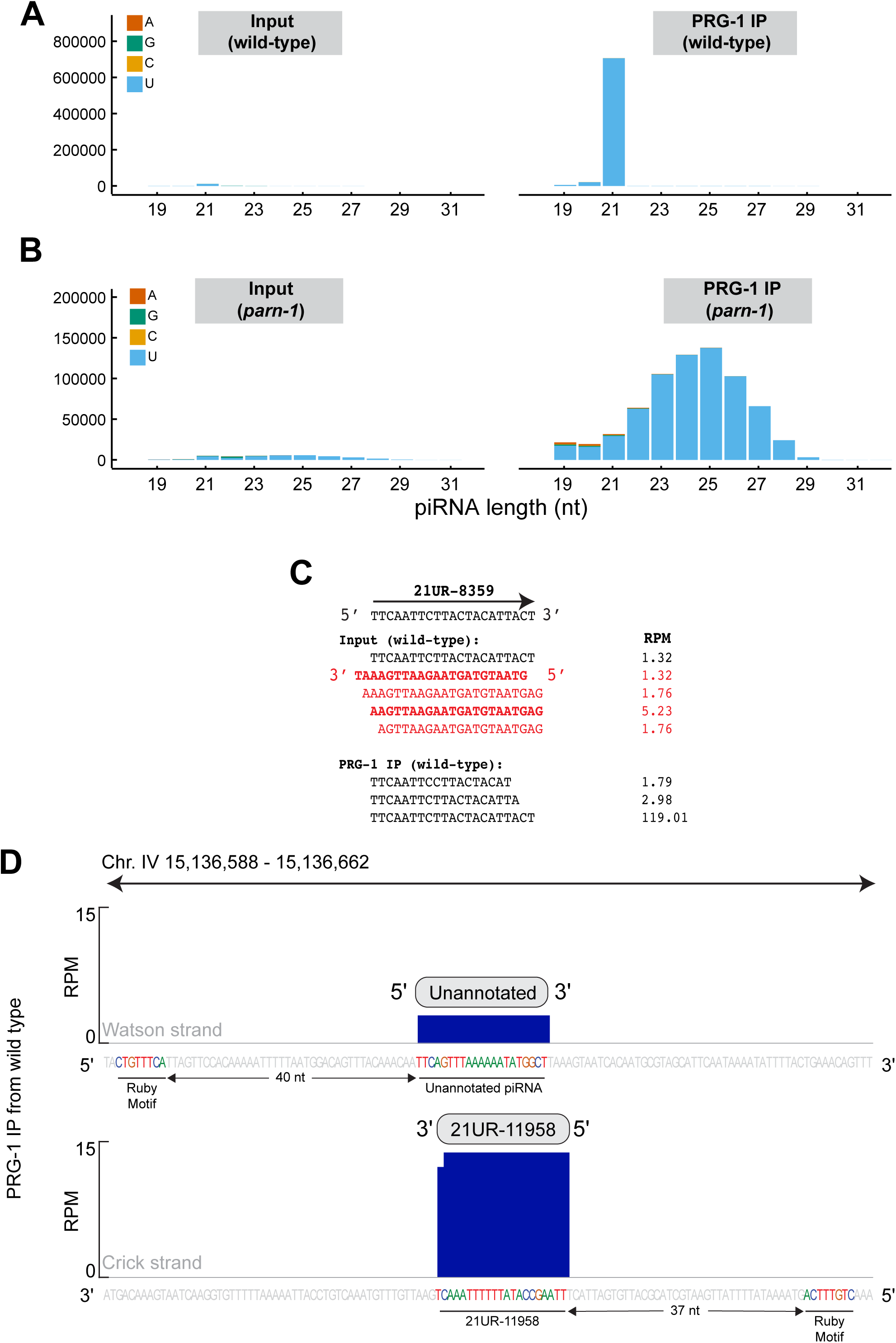
Characters of piRNA and anti-piRNA in wild-type and *parn-1* mutants. Related to Figure 1. A) Barplots showing the RPM of length and first nucleotide of piRNA mapping sequences in wild-type input and PRG-1 IP samples. Data are obtained from one biological replicate. B) Same as (A) but in *parn-1* mutants. C) Sequencing reads mapping to the *21ur-8359* genomic locus from wild-type input and PRG-1 IP. Shown in red are antisense sequences. RPM for individual reads is displayed. D) Genome browser view showing *21ur-11958* locus that overlaps a putative and unannotated piRNA gene. The figure displays a 125 nt window (Chr. IV 15,136,588-15,136,662). The top browser view shows the genomic sequence residing on the Watson strand, highlighted are the Ruby motif and putative piRNA gene. The bottom row shows the genomic sequence residing on the Crick strand. Highlighted are the *21ur-11958* locus and upstream Ruby motif. Blue bars represent the RPM.

**Supplementary Figure S2.**
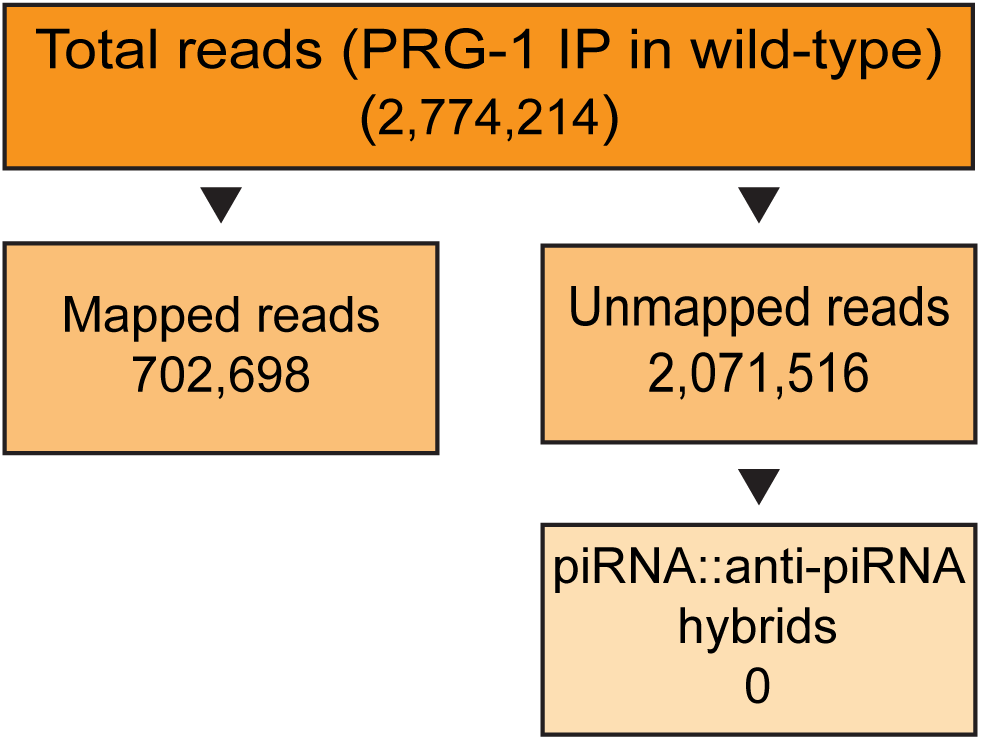
Analysis of PRG-1 IP followed by RNA ligation in wild-type. Related to Figure 2. A) Flow chart showing library statistics and the number of piRNA::anti-piRNA chimeras from PRG-1 IP followed by RNA ligation in wild-type.

**Supplementary Figure S3.**
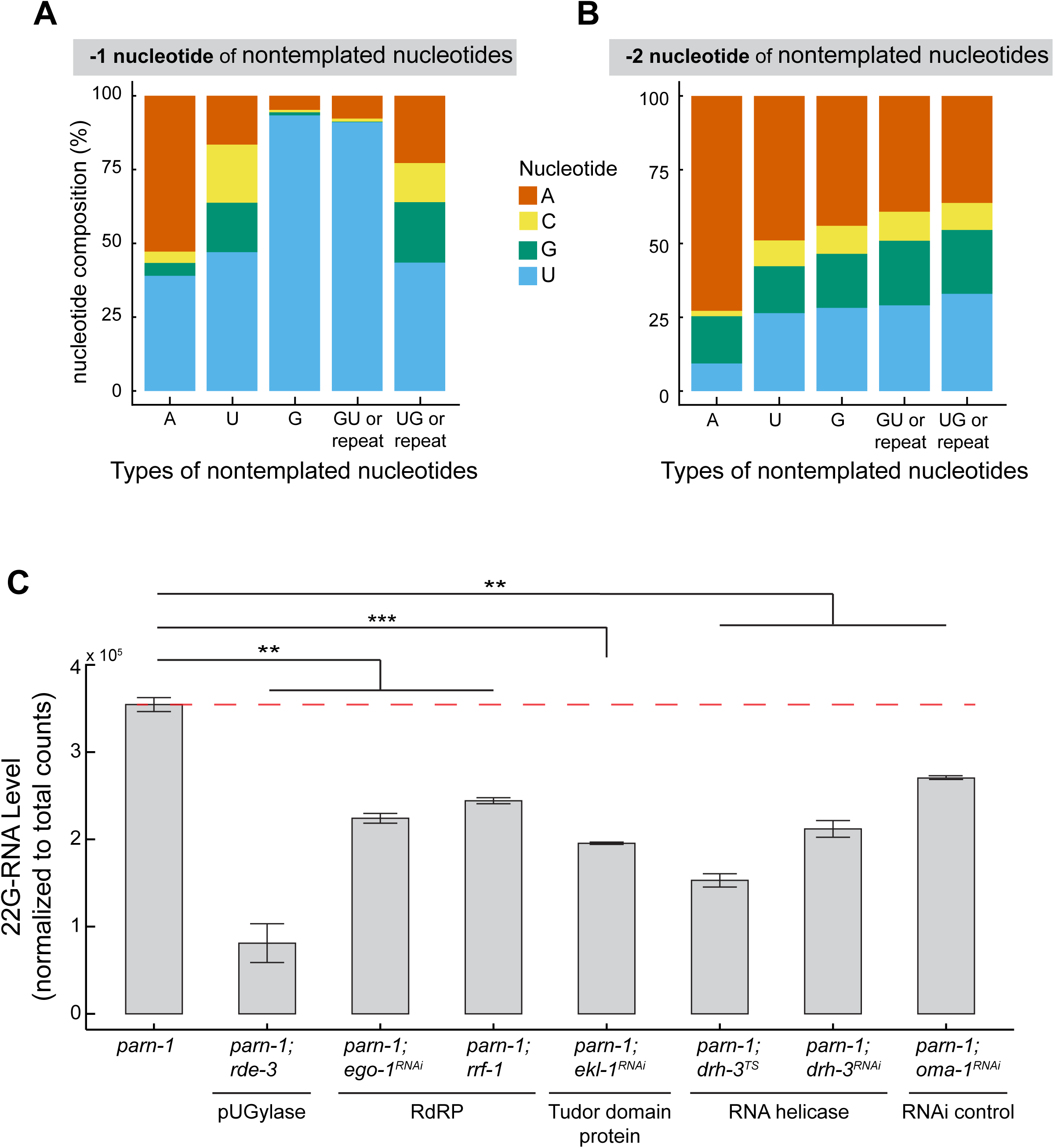
Nucleotide composition of anti-piRNA and 22G-RNA levels. Related to Figure 3. A) Barplots showing the nucleotide composition immediately preceding non-templated A, U, G, GU/GU repeat, and UG/UG repeat tails in *parn-1* mutants. B) Same as (B) but looking at the position two nucleotides upstream of the non-templated nucleotides. C) 22G-RNA levels in *parn-1, parn-1; rde-3, parn-1; ego-1^RNAi^, parn-1; rrf-1, parn-1; ekl-1^RNAi^, parn1; drh-3^TS^, parn-1; drh-3^RNAi^*and *parn-1; oma-1^RNAi^*. Data are displayed as the mean ± standard deviation of 2 biological replicates. p < 0.01 **, p < 0.001 ***, two-sided T-test.

**Supplementary Figure S4.**
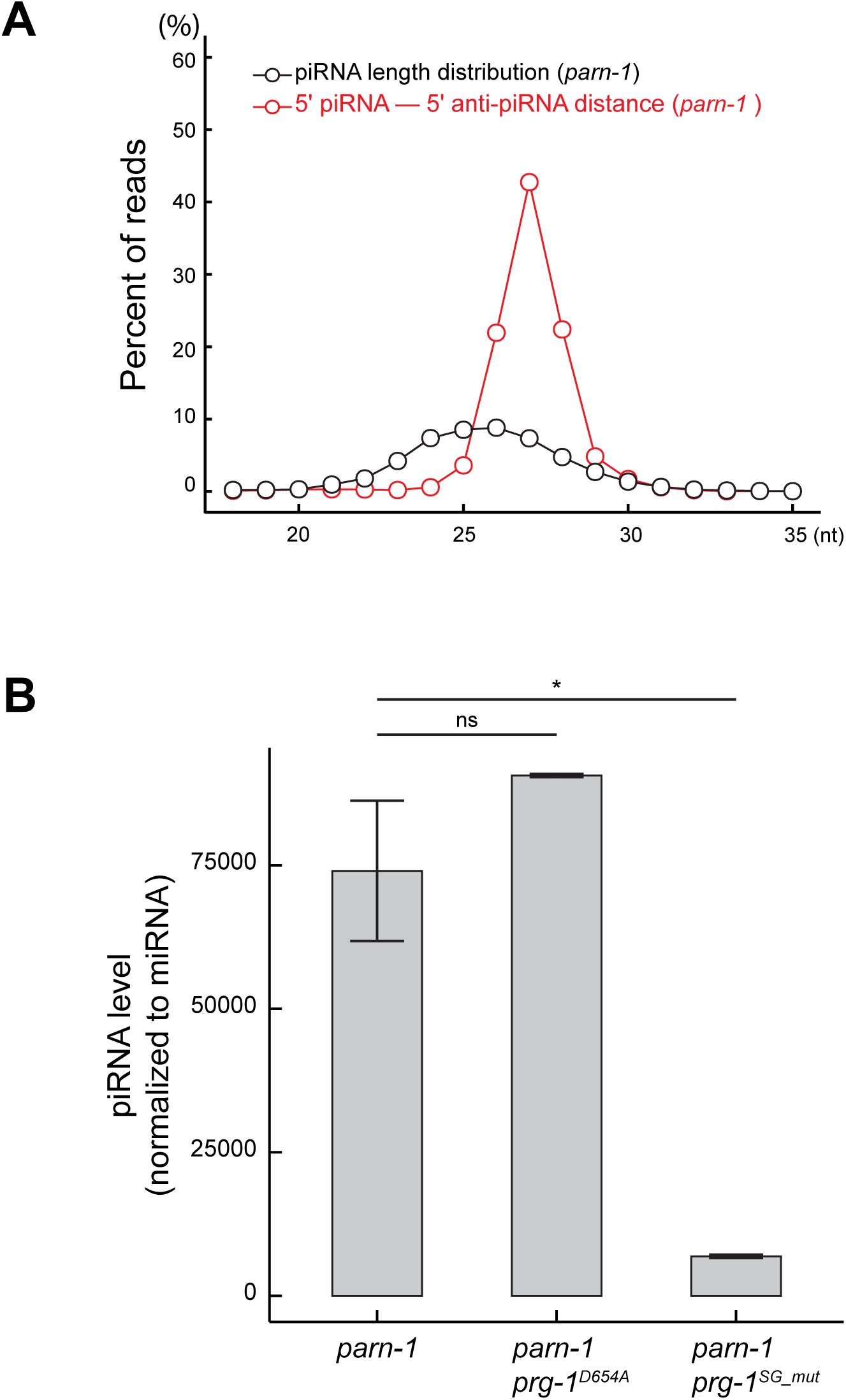
5′ to 5′ distance of piRNAs to anti-piRNAs. Related to Figure 4. A) Line plot showing the length distribution of piRNAs in *parn-1* (black), as well as the 5′ to 5′ distance of anti-piRNAs to piRNAs in *parn-1* mutants (red). B) Total piRNA level in *parn-1, parn-1; prg-1^DA^, and parn-1; prg-1^SG_mut^.* Data are displayed as the mean ± standard deviation of 2 biological replicates. ns: no significance, p < 0.05 *.

**Supplementary Figure S5.**
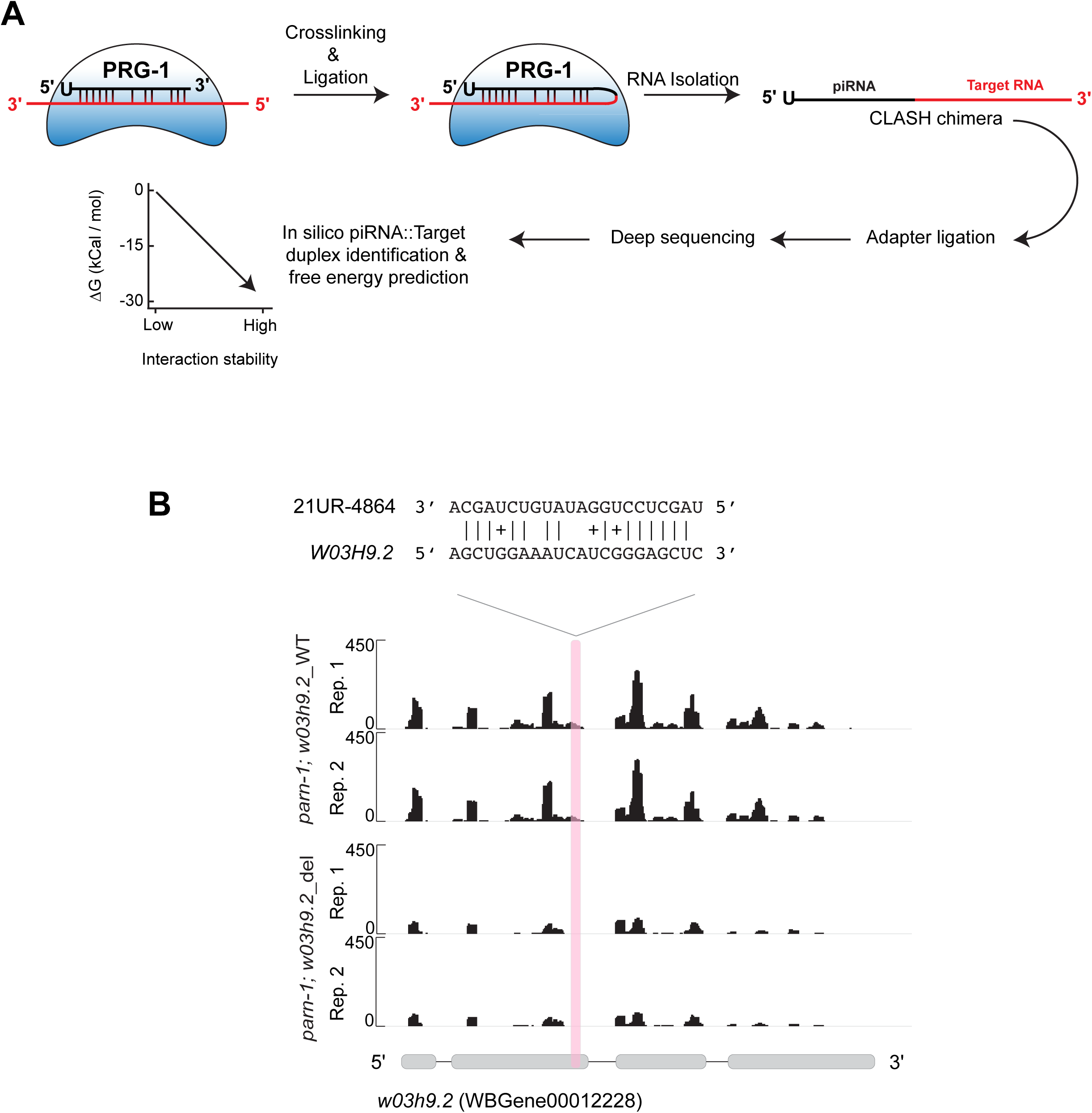
CLASH procedure and 22G-RNA coverage at *w03h9.2*. Related to Figure 5. A) Schematic illustrating the Crosslinking Ligation and Sequencing of Hybrids (CLASH) workflow. piRNAs and targets were crosslinked to PRG-1 protein and subsequently ligated to. yield the CLASH chimera. Chimeras are subjected to adapter ligation and deep sequencing. After sequencing CLASH chimeras can then be identified, piRNA::target duplexes can be predicted and free energy calculation for piRNA::target duplexes can be calculated. The inverse relationship between 1′G and interaction stability is shown. B) Coverage of 22G-RNAs mapping to *w03h9.2*. The precise position of the 21UR-4864 piRNA target site is highlighted as a red bar. Shown above the browser tracts is the interaction between 21UR-4864 and *w03h9.2* transcript as detected by CLASH. Base-pairing between 21UR-4864 and w03h9.2 is indicated with “|”, GU base-pairing is indicated with “+”. Data are displayed as individual biological replicates.

**Supplementary Table S1: Strains and alleles**

**Supplementary Table S2: piRNA abundance in *wild-type*, *parn-1* and *prg-1; parn-1* strains**

**Supplementary Table S3: anti-piRNA abundance in *wild-type* and *parn-1* mutants**

## REFERENCES

1. Ozata, D.M., Gainetdinov, I., Zoch, A., O’Carroll, D. and Zamore, P.D. (2019) PIWI-interacting RNAs: small RNAs with big functions. Nature reviews. Genetics, 20, 89–108.

2. Weick, E.M. and Miska, E.A. (2014) piRNAs: from biogenesis to function. Development, 141, 3458–3471.

3. Luteijn, M.J. and Ketting, R.F. (2013) PIWI-interacting RNAs: from generation to transgenerational epigenetics. Nature reviews. Genetics, 14, 523–534.

4. Wang, X., Ramat, A., Simonelig, M. and Liu, M.F. (2023) Emerging roles and functional mechanisms of PIWI-interacting RNAs. Nature reviews. Molecular cell biology, 24, 123–141.

5. Cox, D.N., Chao, A., Baker, J., Chang, L., Qiao, D. and Lin, H. (1998) A novel class of evolutionarily conserved genes defined by piwi are essential for stem cell self-renewal. Genes & development, 12, 3715–3727.

6. Aravin, A.A., Hannon, G.J. and Brennecke, J. (2007) The Piwi-piRNA pathway provides an adaptive defense in the transposon arms race. Science, 318, 761–764.

7. Malone, C.D. and Hannon, G.J. (2009) Small RNAs as guardians of the genome. Cell, 136, 656–668.

8. Siomi, M.C., Sato, K., Pezic, D. and Aravin, A.A. (2011) PIWI-interacting small RNAs: the vanguard of genome defence. Nature reviews. Molecular cell biology, 12, 246–258.

9. Yu, T., Koppetsch, B.S., Pagliarani, S., Johnston, S., Silverstein, N.J., Luban, J., Chappell, K., Weng, Z. and Theurkauf, W.E. (2019) The piRNA Response to Retroviral Invasion of the Koala Genome. Cell, 179, 632–643 e612.

10. Shirayama, M., Seth, M., Lee, H.C., Gu, W., Ishidate, T., Conte, D., Jr. and Mello, C.C. (2012) piRNAs initiate an epigenetic memory of nonself RNA in the C. elegans germline. Cell, 150, 65–77.

11. Ashe, A., Sapetschnig, A., Weick, E.M., Mitchell, J., Bagijn, M.P., Cording, A.C., Doebley, A.L., Goldstein, L.D., Lehrbach, N.J., Le Pen, J., et al. (2012) piRNAs can trigger a multigenerational epigenetic memory in the germline of C. elegans. Cell, 150, 88–99.

12. Horwich, M.D., Li, C., Matranga, C., Vagin, V., Farley, G., Wang, P. and Zamore, P.D. (2007) The Drosophila RNA methyltransferase, DmHen1, modifies germline piRNAs and single-stranded siRNAs in RISC. Current biology: CB, 17, 1265-1272.

13. Saito, K., Sakaguchi, Y., Suzuki, T., Suzuki, T., Siomi, H. and Siomi, M.C. (2007) Pimet, the Drosophila homolog of HEN1, mediates 2’-O-methylation of Piwi-interacting RNAs at their 3’ ends. Genes & development, 21, 1603–1608.

14. Kirino, Y. and Mourelatos, Z. (2007) Mouse Piwi-interacting RNAs are 2’-O-methylated at their 3’ termini. Nature structural & molecular biology, 14, 347–348.

15. Batista, P.J., Ruby, J.G., Claycomb, J.M., Chiang, R., Fahlgren, N., Kasschau, K.D., Chaves, D.A., Gu, W., Vasale, J.J., Duan, S. et al. (2008) PRG-1 and 21U-RNAs interact to form the piRNA complex required for fertility in C. elegans. Molecular cell, 31, 67–78.

16. Gu, W., Lee, H.C., Chaves, D., Youngman, E.M., Pazour, G.J., Conte, D., Jr. and Mello, C.C. (2012) CapSeq and CIP-TAP identify Pol II start sites and reveal capped small RNAs as C. elegans piRNA precursors. Cell, 151, 1488–1500.

17. Ruby, J.G., Jan, C., Player, C., Axtell, M.J., Lee, W., Nusbaum, C., Ge, H. and Bartel, D.P. (2006) Large-scale sequencing reveals 21U-RNAs and additional microRNAs and endogenous siRNAs in C. elegans. Cell, 127, 1193–1207.

18. Das, P.P., Bagijn, M.P., Goldstein, L.D., Woolford, J.R., Lehrbach, N.J., Sapetschnig, A., Buhecha, H.R., Gilchrist, M.J., Howe, K.L., Stark, R. et al. (2008) Piwi and piRNAs act upstream of an endogenous siRNA pathway to suppress Tc3 transposon mobility in the Caenorhabditis elegans germline. Molecular cell, 31, 79–90.

19. Wang, G. and Reinke, V. (2008) A C. elegans Piwi, PRG-1, regulates 21U-RNAs during spermatogenesis. Current biology: CB, 18, 861-867.

20. Pastore, B., Hertz, H.L. and Tang, W. (2022) Comparative analysis of piRNA sequences, targets and functions in nematodes. RNA biology, 19, 1276–1292.

21. Weick, E.M., Sarkies, P., Silva, N., Chen, R.A., Moss, S.M., Cording, A.C., Ahringer, J., Martinez-Perez, E. and Miska, E.A. (2014) PRDE-1 is a nuclear factor essential for the biogenesis of Ruby motif-dependent piRNAs in C. elegans. Genes & development, 28, 783–796.

22. Kasper, D.M., Wang, G., Gardner, K.E., Johnstone, T.G. and Reinke, V. (2014) The C. elegans SNAPc component SNPC-4 coats piRNA domains and is globally required for piRNA abundance. Developmental cell, 31, 145–158.

23. Goh, W.S., Seah, J.W., Harrison, E.J., Chen, C., Hammell, C.M. and Hannon, G.J. (2014) A genome-wide RNAi screen identifies factors required for distinct stages of C. elegans piRNA biogenesis. Genes & development, 28, 797–807.

24. Weng, C., Kosalka, J., Berkyurek, A.C., Stempor, P., Feng, X., Mao, H., Zeng, C., Li, W.J., Yan, Y.H., Dong, M.Q. et al. (2019) The USTC co-opts an ancient machinery to drive piRNA transcription in C. elegans. Genes & development, 33, 90–102.

25. Podvalnaya, N., Bronkhorst, A.W., Lichtenberger, R., Hellmann, S., Nischwitz, E., Falk, T., Karaulanov, E., Butter, F., Falk, S. and Ketting, R.F. (2023) piRNA processing by a trimeric Schlafen-domain nuclease. bioRxiv, 2023.2001.2019.524756.

26. Kamminga, L.M., van Wolfswinkel, J.C., Luteijn, M.J., Kaaij, L.J., Bagijn, M.P., Sapetschnig, A., Miska, E.A., Berezikov, E. and Ketting, R.F. (2012) Differential impact of the HEN1 homolog HENN-1 on 21U and 26G RNAs in the germline of Caenorhabditis elegans. PLoS genetics, 8, e1002702.

27. Montgomery, T.A., Rim, Y.S., Zhang, C., Dowen, R.H., Phillips, C.M., Fischer, S.E. and Ruvkun, G. (2012) PIWI associated siRNAs and piRNAs specifically require the Caenorhabditis elegans HEN1 ortholog henn-1. PLoS genetics, 8, e1002616.

28. Tang, W., Tu, S., Lee, H.C., Weng, Z. and Mello, C.C. (2016) The RNase PARN-1 Trims piRNA 3’ Ends to Promote Transcriptome Surveillance in C. elegans. Cell, 164, 974–984.

29. Billi, A.C., Alessi, A.F., Khivansara, V., Han, T., Freeberg, M., Mitani, S. and Kim, J.K. (2012) The Caenorhabditis elegans HEN1 ortholog, HENN-1, methylates and stabilizes select subclasses of germline small RNAs. PLoS Genet., 8, e1002617.

30. Izumi, N., Shoji, K., Sakaguchi, Y., Honda, S., Kirino, Y., Suzuki, T., Katsuma, S. and Tomari, Y. (2016) Identification and Functional Analysis of the Pre-piRNA 3’ Trimmer in Silkworms. Cell, 164, 962–973.

31. Ding, D., Liu, J., Dong, K., Midic, U., Hess, R.A., Xie, H., Demireva, E.Y. and Chen, C. (2017) PNLDC1 is essential for piRNA 3’ end trimming and transposon silencing during spermatogenesis in mice. Nature communications, 8, 819.

32. Zhang, Y., Guo, R., Cui, Y., Zhu, Z., Zhang, Y., Wu, H., Zheng, B., Yue, Q., Bai, S., Zeng, W. et al. (2017) An essential role for PNLDC1 in piRNA 3’ end trimming and male fertility in mice. Cell research.

33. Nishimura, T., Nagamori, I., Nakatani, T., Izumi, N., Tomari, Y., Kuramochi-Miyagawa, S. and Nakano, T. (2018) PNLDC1, mouse pre-piRNA Trimmer, is required for meiotic and post-meiotic male germ cell development. EMBO reports, 19.

34. Nagirnaja, L., Morup, N., Nielsen, J.E., Stakaitis, R., Golubickaite, I., Oud, M.S., Winge, S.B., Carvalho, F., Aston, K.I., Khani, F. et al. (2021) Variant PNLDC1, Defective piRNA Processing, and Azoospermia. N Engl J Med, 385, 707–719.

35. Wang, X., Tan, Y.Q. and Liu, M.F. (2022) Defective piRNA Processing and Azoospermia. N Engl J Med, 386, 1674–1675.

36. Lee, H.C., Gu, W., Shirayama, M., Youngman, E., Conte, D., Jr. and Mello, C.C. (2012) C. elegans piRNAs mediate the genome-wide surveillance of germline transcripts. Cell, 150, 78–87.

37. Bagijn, M.P., Goldstein, L.D., Sapetschnig, A., Weick, E.M., Bouasker, S., Lehrbach, N.J., Simard, M.J. and Miska, E.A. (2012) Function, targets, and evolution of Caenorhabditis elegans piRNAs. Science, 337, 574–578.

38. Buckley, B.A., Burkhart, K.B., Gu, S.G., Spracklin, G., Kershner, A., Fritz, H., Kimble, J., Fire, A. and Kennedy, S. (2012) A nuclear Argonaute promotes multigenerational epigenetic inheritance and germline immortality. Nature, 489, 447–451.

39. Smardon, A., Spoerke, J.M., Stacey, S.C., Klein, M.E., Mackin, N. and Maine, E.M. (2000) EGO-1 is related to RNA-directed RNA polymerase and functions in germ-line development and RNA interference in C. elegans. Current biology: CB, 10, 169–178.

40. Gu, W., Shirayama, M., Conte, D., Jr., Vasale, J., Batista, P.J., Claycomb, J.M., Moresco, J.J., Youngman, E.M., Keys, J., Stoltz, M.J. et al. (2009) Distinct argonaute-mediated 22G-RNA pathways direct genome surveillance in the C. elegans germline. Molecular cell, 36, 231–244.

41. Sijen, T., Fleenor, J., Simmer, F., Thijssen, K.L., Parrish, S., Timmons, L., Plasterk, R.H. and Fire, A. (2001) On the role of RNA amplification in dsRNA-triggered gene silencing. Cell, 107, 465–476.

42. Yigit, E., Batista, P.J., Bei, Y., Pang, K.M., Chen, C.C., Tolia, N.H., Joshua-Tor, L., Mitani, S., Simard, M.J. and Mello, C.C. (2006) Analysis of the C. elegans Argonaute family reveals that distinct Argonautes act sequentially during RNAi. Cell, 127, 747–757.

43. Tsai, H.Y., Chen, C.C., Conte, D., Jr., Moresco, J.J., Chaves, D.A., Mitani, S., Yates, J.R., 3rd, Tsai, M.D. and Mello, C.C. (2015) A ribonuclease coordinates siRNA amplification and mRNA cleavage during RNAi. Cell, 160, 407-419.

44. Chen, C.C., Simard, M.J., Tabara, H., Brownell, D.R., McCollough, J.A. and Mello, C.C. (2005) A member of the polymerase beta nucleotidyltransferase superfamily is required for RNA interference in C. elegans. Current biology: CB, 15, 378–383.

45. Shukla, A., Yan, J., Pagano, D.J., Dodson, A.E., Fei, Y., Gorham, J., Seidman, J.G., Wickens, M. and Kennedy, S. (2020) poly(UG)-tailed RNAs in genome protection and epigenetic inheritance. Nature, 582, 283–288.

46. Preston, M.A., Porter, D.F., Chen, F., Buter, N., Lapointe, C.P., Keles, S., Kimble, J. and Wickens, M. (2019) Unbiased screen of RNA tailing activities reveals a poly(UG) polymerase. Nature methods, 16, 437–445.

47. Shukla, A., Perales, R. and Kennedy, S. (2021) piRNAs coordinate poly(UG) tailing to prevent aberrant and perpetual gene silencing. Current biology: CB, 31, 4473–4485 e4473.

48. Ghanta, K.S., Ishidate, T. and Mello, C.C. (2021) Microinjection for precision genome editing in Caenorhabditis elegans. STAR Protoc, 2, 100748.

49. Qu, W., Ren, C., Li, Y., Shi, J., Zhang, J., Wang, X., Hang, X., Lu, Y., Zhao, D. and Zhang, C. (2011) Reliability analysis of the Ahringer Caenorhabditis elegans RNAi feeding library: a guide for genome-wide screens. BMC Genomics, 12, 170.

50. Li, L., Dai, H., Nguyen, A.P. and Gu, W. (2020) A convenient strategy to clone small RNA and mRNA for high-throughput sequencing. Rna, 26, 218–227.

51. Martin, M. (2011) Cutadapt removes adapter sequences from high-throughput sequencing reads. 2011, 17, 3.

52. Langmead, B., Trapnell, C., Pop, M. and Salzberg, S.L. (2009) Ultrafast and memory-efficient alignment of short DNA sequences to the human genome. Genome Biology, 10, R25.

53. Neph, S., Kuehn, M.S., Reynolds, A.P., Haugen, E., Thurman, R.E., Johnson, A.K., Rynes, E., Maurano, M.T., Vierstra, J., Thomas, S. et al. (2012) BEDOPS: high-performance genomic feature operations. Bioinformatics, 28, 1919–1920.

54. Quinlan, A.R. and Hall, I.M. (2010) BEDTools: a flexible suite of utilities for comparing genomic features. Bioinformatics, 26, 841–842.

55. Kent, W.J., Sugnet, C.W., Furey, T.S., Roskin, K.M., Pringle, T.H., Zahler, A.M., Haussler and David. (2002) The Human Genome Browser at UCSC. Genome Research, 12, 996–1006.

56. Robinson, J.T., Thorvaldsdóttir, H., Winckler, W., Guttman, M., Lander, E.S., Getz, G. and Mesirov, J.P. (2011) Integrative genomics viewer. Nature Biotechnology, 29, 24–26.

57. Chou, M.T., Han, B.W., Hsiao, C.P., Zamore, P.D., Weng, Z. and Hung, J.H. (2015) Tailor: a computational framework for detecting non-templated tailing of small silencing RNAs. Nucleic acids research, 43, e109.

58. Shen, E.Z., Chen, H., Ozturk, A.R., Tu, S., Shirayama, M., Tang, W., Ding, Y.H., Dai, S.Y., Weng, Z. and Mello, C.C. (2018) Identification of piRNA Binding Sites Reveals the Argonaute Regulatory Landscape of the C. elegans Germline. Cell, 172, 937–951 e918.

59. Gruber, A.R., Lorenz, R., Bernhart, S.H., Neuböck, R. and Hofacker, I.L. (2008) The Vienna RNA websuite. Nucleic Acids Res, 36, W70–74.

60. Pastore, B., Hertz, H.L., Price, I.F. and Tang, W. (2021) pre-piRNA trimming and 2’-O-methylation protect piRNAs from 3’ tailing and degradation in C. elegans. Cell reports, 36, 109640.

61. Seroussi, U., Lugowski, A., Wadi, L., Lao, R.X., Willis, A.R., Zhao, W., Sundby, A.E., Charlesworth, A.G., Reinke, A.W. and Claycomb, J.M. (2023) A comprehensive survey of C. elegans argonaute proteins reveals organism-wide gene regulatory networks and functions. eLife, 12.

62. Carninci, P., Sandelin, A., Lenhard, B., Katayama, S., Shimokawa, K., Ponjavic, J., Semple, C.A., Taylor, M.S., Engstrom, P.G., Frith, M.C. et al. (2006) Genome-wide analysis of mammalian promoter architecture and evolution. Nat Genet, 38, 626–635.

63. Carninci, P., Kasukawa, T., Katayama, S., Gough, J., Frith, M.C., Maeda, N., Oyama, R., Ravasi, T., Lenhard, B., Wells, C. et al. (2005) The transcriptional landscape of the mammalian genome. Science, 309, 1559–1563.

64. Imburgio, D., Rong, M., Ma, K. and McAllister, W.T. (2000) Studies of promoter recognition and start site selection by T7 RNA polymerase using a comprehensive collection of promoter variants. Biochemistry, 39, 10419–10430.

65. Saito, T.L., Hashimoto, S., Gu, S.G., Morton, J.J., Stadler, M., Blumenthal, T., Fire, A. and Morishita, S. (2013) The transcription start site landscape of C. elegans. Genome research, 23, 1348–1361.

66. Claycomb, J.M., Batista, P.J., Pang, K.M., Gu, W., Vasale, J.J., van Wolfswinkel, J.C., Chaves, D.A., Shirayama, M., Mitani, S., Ketting, R.F., et al. (2009) The Argonaute CSR-1 and its 22G-RNA cofactors are required for holocentric chromosome segregation. Cell, 139, 123–134.

67. Kumsta, C. and Hansen, M. (2012) C. elegans rrf-1 mutations maintain RNAi efficiency in the soma in addition to the germline. PloS one, 7, e35428.

68. Jumper, J., Evans, R., Pritzel, A., Green, T., Figurnov, M., Ronneberger, O., Tunyasuvunakool, K., Bates, R., Zidek, A., Potapenko, A. et al. (2021) Highly accurate protein structure prediction with AlphaFold. Nature, 596, 583–589.

69. Anzelon, T.A., Chowdhury, S., Hughes, S.M., Xiao, Y., Lander, G.C. and MacRae, I.J. (2021) Structural basis for piRNA targeting. Nature, 597, 285–289.

70. Helwak, A., Kudla, G., Dudnakova, T. and Tollervey, D. (2013) Mapping the human miRNA interactome by CLASH reveals frequent noncanonical binding. Cell, 153, 654–665.

71. Roschdi, S., Yan, J., Nomura, Y., Escobar, C.A., Petersen, R.J., Bingman, C.A., Tonelli, M., Vivek, R., Montemayor, E.J., Wickens, M. et al. (2022) An atypical RNA quadruplex marks RNAs as vectors for gene silencing. Nature structural & molecular biology, 29, 1113–1121.

